# Cell Type Hierarchy Reconstruction via Reconciliation of Multi-resolution Cluster Tree

**DOI:** 10.1101/2021.02.06.430067

**Authors:** Minshi Peng, Brie Wamsley, Andrew Elkins, Daniel M Geschwind, Yuting Wei, Kathryn Roeder

## Abstract

A wealth of clustering algorithms are available for Single-cell RNA sequencing (scRNA-seq), but it remains challenging to compare and characterize the features across different scales of resolution. To resolve this challenge Multi-resolution Reconciled Tree (MRtree), builds a hierarchical tree structure based on multi-resolution partitions that is highly flexible and can be coupled with most scRNA-seq clustering algorithms. MRtree out-performs bottom-up or divisive hierarchical clustering approaches because it inherits the robustness and versatility of a flat clustering approach, while maintaining the hierarchical structure of cells. Application to fetal brain cells yields insight into subtypes of cells that can be reliably estimated.

## Background

Single-cell RNA sequencing (scRNA-seq) is a recently developed technology that is being widely deployed to collect unprecedented catalogues detailing the transcriptomes of individual cells. The ability to capture the molecular heterogeneity of tissues at high resolution underlies its increasing popularity in discovering cellular and molecular underpinnings of complex and rare cell populations^2^, developing a “parts list” for complex tissues ^13,20^, and studying various diseases and cell development or lineage. One essential component involves the utility of scRNA-seq data to enable the identification of functionally distinct subpopulations that each possess a different pattern of gene expression activity. These sub-populations can indicate different cell types with relatively stable, static behavior, or cell states in intermediate stages of a transient process. Unbiased discovery of cell types from scRNA-seq data can be automated using a wealth of unsupervised clustering algorithms, among them the most widely-applied include the Louvain graph-based algorithm incorporated as part of Seurat ^3,10^ pipeline, k-means clustering and its derivatives by consensus clustering as performed in SC3^7^ and SIMLR^12^, and other methods that address issues caused by rare types such as RaceID ^4^.

A major challenge regarding clustering algorithms mentioned above is that they explicitly or implicitly require the number of clusters to be supplied as an input parameter. Determining the number of cell types in the population presents a significant challenge given the large number of distinct cell types, which is further complicated by substantial biological and technical variation. There are some computational methods available to guide the choice of *K*; however, these methods are shown to be flawed in one aspect or another. For example, methods developed on the basis of statistical testing are shown to be too sensitive to heterogeneity, especially for large samples, while other methods tend to favor a fairly coarse resolution, with clearly separated clusters and fail to identify closely related and overlapping cell types. Therefore, judgment from the researchers is required to choose the desired resolution. A common practice in scRNA-seq data analysis is to run a clustering algorithm repeatedly for a range of resolutions, followed by careful inspections of individual results by examining the cluster compositions and the expression of published marker genes to select the final partition. This supervised process takes significant time and effort and is limited by the current state of the investigator’s/field’s knowledge about cell type and cell state diversity. It would be a substantial advantage in terms of efficiency and veracity to be able to reach the same level of resolution in an unsupervised manner.

Hierarchical clustering (HC) is another popular general-purpose clustering method commonly used to identify cell-populations ^18,1,16^. HC has the advantage of being able to determine relationships between clusters of different granularities since the result can be visualized as a dendrogram. This dendrogram is then “cut” at different heights to generate different numbers of clusters. This hierarchical structure helps identify multiple levels of functional specialization of cells. For instance, neurons share specific functional characteristics distinct from those of various glial cell types and contain distinct subtypes with more specialized functions, such as excitatory or inhibitory properties. Different variants of hierarchical clustering make different assumptions, the most common ones used in classical hierarchical agglomerative clustering (HAC) is Ward’s ^15^ and “average” linkage as adopted in SC3^7^. An important limitation of HAC is that both time and memory requirements scale at least quadratically with the number of data points, which is slower than many flat clustering methods like K-means and prohibitively expensive for large data sets. A few scRNA-seq tools expand upon the idea of hierarchical clustering, for instance, pcaReduce^22^ introduces an agglomerative clustering approach by conducting dimension reduction after each merge, starting from an initial clustering, and CellBIC ^6^ performs bisecting clustering in a top-down manner leveraging the bi-modal gene expression patterns. However, these methods either require a good initial start that is implicitly equivalent to the choice of *K*, or are highly dependent on the assumption of bimodal expression pattern at each iteration, which is not appropriate for multi-modal data.

In this study, we build a tool to bridge the gap between two separate lines of inquiry, flat and hierarchical clustering. Empirically, scRNA-seq data analysts observe that the partitions obtained from flat clustering at multiple resolutions, when ordered by increasing resolution, produce a layered structure with a tree-style backbone ^17^. This produces a useful representation to help visually determine the stability of clusters and relations among them. We build on this idea and propose a method called Multi-resolution Reconciled Tree (MRtree) that reconstructs the underlying tree structure by reconciling partitions obtained at different granularities (Figure 1) to produce a coherent hierarchy that is as similar as possible to the original flat clustering at different scales. It can work with many specially-designed flat clustering algorithms for single-cell data, such as Louvain clustering from Seurat ^3^, thus inheriting the scalability and good performance in clustering the single-cell data; meanwhile, it recovers the intrinsic hierarchy structure determined by the cell types and cell states.

**Figure 1:**
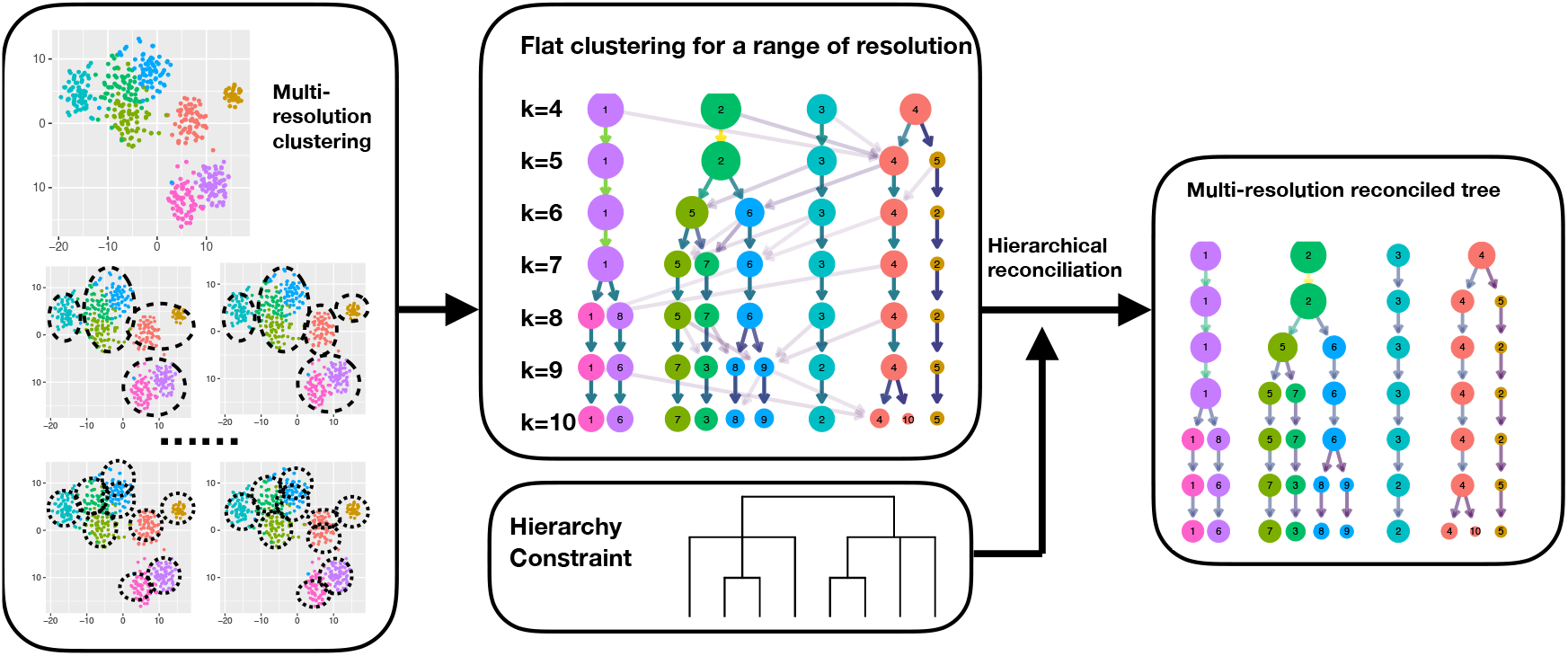
Overview of MRtree framework. The algorithm starts by performing flat clustering on scRNA-seq data for a range of resolutions, where the partitions between adjacent resolutions are matched to form a graph as an entangled cluster tree. Then reconciliation is performed through optimization with the hierarchical structure enforced by constraints. The obtained final optimal solution represents the recovered hierarchical cluster tree.

Applications of MRtree on a variety of scRNA-seq data sets, including mouse brain^18^, human pancreas^14,8^ and human fetal brain ^9^, showed improved performance for clustering of scRNA-seq data over initial flat clustering methods. The hierarchical structure discovered by MRtree easily outperformed a variety of tree-construction methods. Moreover, the results accurately reflect the extent of transcriptional distinctions among cell groups and align well with levels of functional specializations among cells. Particularly, when applied to developing human brain cells, the method successfully identified major cell types and recovered an underlying hierarchical structure that is highly consistent with the results from the original study ^9^. Subsequent analysis on each major type via MRtree revealed finer sub-structure defining biologically plausible subtypes, determined mainly by maturation states, spatial location, and terminal specification.

## Results

### Methods overview

MRtree aims to recover a hierarchical tree by denoising and integrating a series of flat clusterings into a coherent tree structure. The algorithm starts by applying a suitable flat clustering algorithm to obtain partitions for a range of resolution parameters. The multi-scale results can be represented using a multi-partite graph, referred to as a *cluster tree*, where the nodes represent clusters, and edges between partitions of adjacent resolutions indicate common cells shared. We propose an efficient optimization procedure to reconcile the incoherent cell assignments across resolutions, that produces the optimal underlying tree structure following the hierarchy constraints, while adhering to the initial flat clustering to the maximum extent. Formally, this is achieved by minimizing (among valid hierarchical tree structures), the difference between initial multi-level cluster assignments and the cluster assignments in the resulting tree structure. By representing the partitions as a multi-partite graph, the clustering assignments that violate the hierarchy constraint can be identified as merging directed edges and thus penalized in the objective function. The optimization procedure proceeds by iteratively and greedily identifying those tree nodes, which, when corrected by reassigning the associated conflicting cell lineages, contribute to maximum descent in the defined objective function. The outcome of the proposed optimization procedure is a reconciled tree, named the *hierarchical cluster tree*, representing the optimal tree-based cluster arrangement across scales (Figure 1, Figure S1, Supplemental Information).

Our method is motivated by consensus clustering (also known as ensemble clustering); however, instead of gathering information over repeated runs of algorithms at the same resolution, we leverage the cluster structure revealed at multiple scales to build an ensembled hierarchy. The common features across resolutions are identified and averaged to reduce noise, while the distinctions between resolutions are utilized to uncover different scales of geometric structure, which are further reconciled to conform to a robust hierarchical tree. We stress that consensus clustering is essentially a noise-reduction technique that aims to deliver robust, interpretable results.

Another key distinguishing feature of our procedure, compared to existing consensus methods, is that we build a cluster hierarchy directly from the flat partitions in an “in place” way. This is in comparison to existing methods for which an alternative hierarchical clustering algorithm is applied to the co-classification consensus matrix built from an ensemble of partitions. MRtree uses an optimization framework to edit the original partitions through a similar voting scheme. At the same time, it aims to preserve the original splitting order of the hierarchy determined by the clustering algorithm. The proposed method is efficient in terms of memory cost and time complexity (Supplemental Information). Moreover, MRtree enables a direct comparison of partitions before and after tree reconciliation, to examine the stability of the clustering algorithm at different scales. As a benefit, we are able to trim the tree to the maximum depth within the stable range to obtain reliable final clusters.

To summarize, we present a computationally efficient method to generate a hierarchical tree from flat clustering results for a range of resolutions, by an iterative greedy optimization scheme. It enables us to take advantage of various flat clustering approaches, while improving over these flat clustering results by averaging over membership assignments across resolutions. The resulting hierarchical structure captures the relations between cell types and at the same time helps circumvent the problems of choosing the optimal resolution parameter. It turns out that the proposed method improves the clustering accuracy over the initial partition across scales, and outperforms a variety of alternative tree construction methods for recovering the underlying tree structure. To facilitate identifying stable tree layers, we propose a stability measure that compares the initial flat clustering with the reconciled tree. We also implement tools to sample implicit resolution parameters for Seurat clustering that enable equal coverage of different clustering granularities. Finally, our optimization procedure is made efficient for potential use in big data analyses.

### Simulation study

#### Simulated data

To evaluate how well MRtree is able to recover the cluster hierarchy and improve the clustering across resolutions, we harness the tools provided by the SymSim package^19^ to simulate scRNA-seq data given a known tree structure, using the SymSim parameters estimated from a UMI-based dataset of 3,005 mouse cortex cells ^18^ (Supplemental Information). Motivated by major cell types identified in brain tissues, we constructed a hypothetical tree (Figure 2A,B) as the ground truth representing the hierarchy of the cell types/states.

**Figure 2:**
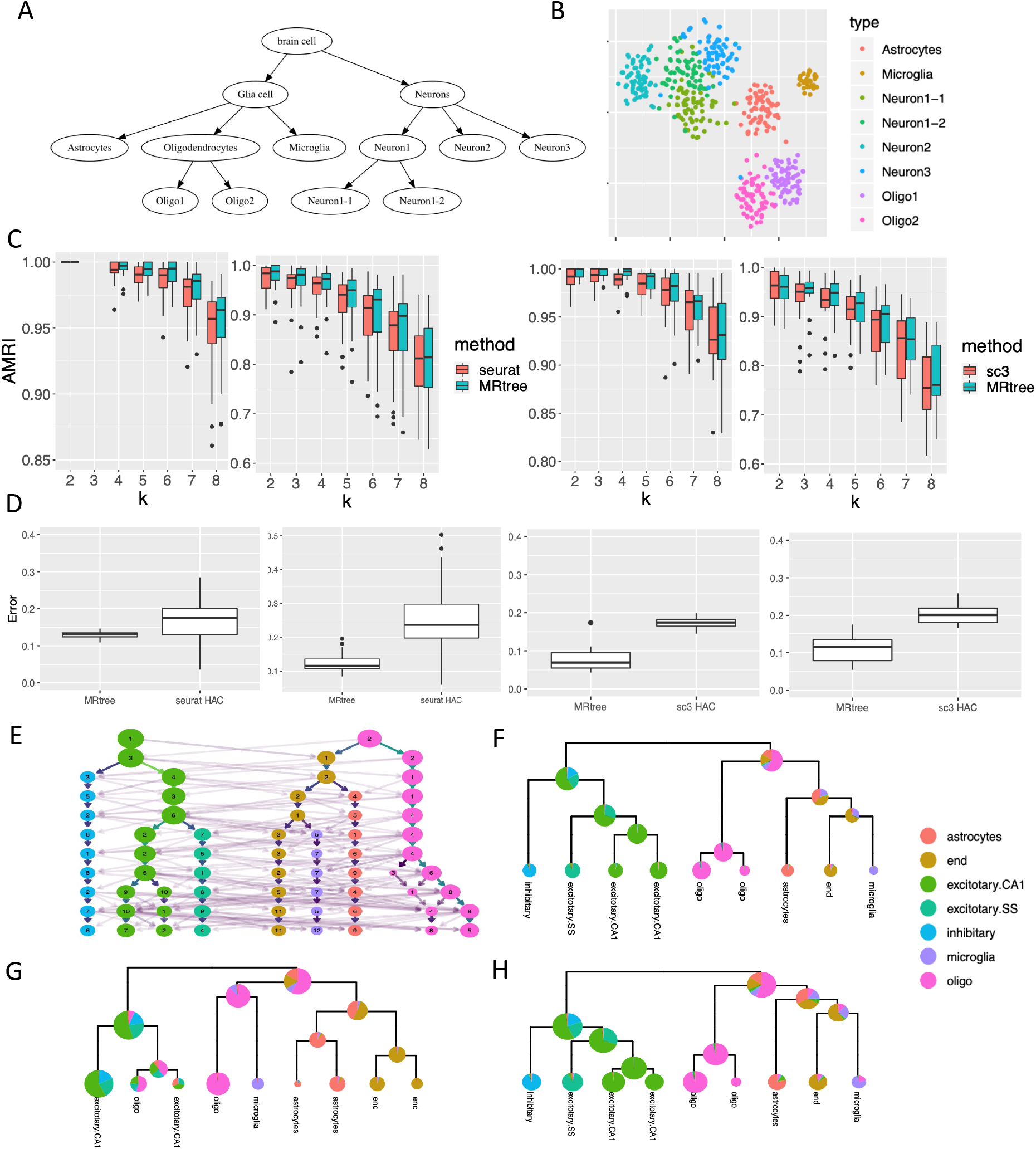
Evaluate the performance of MRtree via simulations and analysis on mouse brain data ^18^. (A) The hypothetical tree structure of the cell states from which cells are generated. (B) tSNE plot of the simulated cells in one experiment, colored by the cell types indicated in the leaf of the actual hierarchical cluster tree. The difficulty of the simulation varies from simple (left, smaller within-cluster noise) to challenging (right, stronger noise) in panels (C) and (D) for each method. (C) Comparing the accuracy of MRtree clusters with the clusters from initial flat clustering at multiple resolutions using Seurat and SC3. The accuracy is measured by the Adjusted Multi-resolution Rand Index (AMRI). (D) Evaluate tree construction accuracy of MRtree with dendrogram from hierarchical clustering obtained with Seurat and SC3. (E-H) MRtree applied to scRNA-seq data from the mouse brain. (E) Initial flat clustering by SOUP on 3,005 cells ^18^ by varying *K*, the resolution parameter specifying the number of clusters, colored by the gold standard labels. (F) The MRtree-constructed tree from initial SOUP clusterings. The pie charts on tree nodes represent the cell type composition referencing the gold standard. (G, H) Comparing tree construction and clustering accuracy on mouse brain data using different methods, hierarchical cluster tree generated by HAC starting with SOUP clusters (G) and starting with individual cells (H).

Repeated simulations were performed by first generating single-cell data with SymSim from the hypothetical tree structure, followed by multi-resolution flat clustering using a variety of clustering methods. Then MRtree was applied to form the hierarchical cluster tree that reconciled the multilevel clusterings. MRtree can be coupled with most flat clustering methods; hence we evaluated the performance using a variety of algorithms, including Seurat ^3^, SC3^7^, SOUP ^21^, and K-means applied to a UMAP projection. The clustering results were evaluated and compared with the raw clusters obtained from flat clustering in three aspects: the accuracy of clustering regarding label assignments at different resolutions, the tree structure estimation accuracy, and the clustering stability.

We first sought to quantify how well MRtree performs regarding clustering accuracy, measured using Adjusted Multiresolution Rand Index (AMRI, Methods) between the obtained labels and true labels known from the simulation. An AMRI close to 1 indicates perfect clustering given the resolution. MRtree achieved higher accuracy almost uniformly across resolutions and for various clustering methods (Figures 2C and S2). It is worth noticing that the reconciliation procedure even improved upon SC3 results, which already employed an ensemble-based method for each fixed resolution. This demonstrates that applying an ensemble approach across resolutions captures additional structural information within the data. In addition, the gain was more pronounced for coarse clustering and when there was more room for improvement.

Next, we evaluated the ability of MRtree to recover the tree structure. For comparison, we leveraged the tools that build hierarchical trees in Seurat and SC3. For Seurat, an agglomerative hierarchical cluster tree was built starting with the identified Seurat clusters, while for SC3, a full HAC was performed from the consensus similarity matrix constructed by aggregating clustering results with different dimension reduction schemes. MRtree produced a significantly improved tree structure estimation compared to the competing methods, as demonstrated by the reduced error of tree reconstruction (Figure 2D, Methods).

Finally we evaluate the clustering stability before and after tree reconciliation, coupled with multiple clustering methods. The stability score is calculated following the subsample procedure described in Methods. For *K* less than the true number of clusters, the measured stability is confounded by the instability induced by the incorrect resolution. Therefore we restrict our comparison to the measured stability at the true resolution. Clustering stability with MRtree is clearly improved compared to the initial clustering across all methods (Figure S3), demonstrating the improved robustness of MRtree, which successfully employs the consensus mechanism to denoise the individual clustering with collective information across resolutions.

### scRNA-seq data

#### Mouse brain cells

We illustrate MRtree using a scRNA-seq data set containing 3,005 cells of somatosensory cortex and hippocampal-CA1 region from mice, collected between postnatal 21-31 days. We call this the mouse brain data^18^. The authors have assigned the cells to seven major types: pyramidal CA1, pyramidal SS, interneurons, astrocytes-ependymal, microglia, endothelial-mural, and oligodendrocytes. For comparison, these labels are treated as the gold standard in the following analysis.

We chose SOUP ^21^ for multi-resolution clustering due to its superior performance on these data. The hard clustering labels were obtained by varying *K*, the resolution parameter specifying the desired number of clusters, from 2 to 12. With MRtree, we were able to construct a hierarchical cluster tree from the flat sequential clusterings. The initial cluster tree is visualized with nodes colored by the major type referencing the gold standard, and the recovered tree from MRtree is shown on the right, with the proportion of cell types in each node visualized by a pie chart (Figure 2E,F). The tree successfully split the neurons and glial cells at an early stage, followed by splitting pyramidal cells from two regions (CA1,SS) from the interneurons. Finally, cells from the same type but distinct brain regions were identified. The tree reconciliation step also improved the clustering performance by increasing the accuracy measured by AMRI in multiple layers (Figure S4a).

To further compare the performance of MRtree with HAC, we applied HAC using complete linkage on the first 20 principal components, starting either form singletons (individual cell) or the 9 SOUP clusters obtained at the maximum resolution (Figure 2G,H; only the top layers of HAC from singleton are shown for comparison purposes). Compared to MRtree, HAC shared a similar overall tree structure, but it generated clusters at lower accuracy for each layer. The results support the argument that MRtree is able to improve accuracy upon initial clustering by pooling information across resolutions. HAC from singletons performed much worse regarding both accuracy and tree structure, possibly owing to the sensitivity of HAC to outliers and linkage selection. For completeness, we also demonstrate the accuracy from two widely applied clustering methods, Seurat and SC3, where the HAC results were generated from the built-in functions provided as part of the toolkits. In both cases, MRtree outperformed both the initial flat clustering and the HAC (Figure S4b,c).

In addition to improving the clustering accuracy, we were able to infer the resolution that achieved the highest stability by inspecting the difference between the initial tree and the reconstructed tree. It indicated that both the SOUP and Seurat algorithms should stop splitting at *K* = 7, which was consistent with the gold standard (Figure S6). Stability analysis on SC3 results showed a preferred resolution of 6 clusters. Indeed, we observed steep drop in accuracy for any resolution greater than 6 (Figure S4c). By comparison, using available *K*-selection methods supported in multiple single-cell analysis pipelines, the optimal number of clusters selected varied widely (Table S1). For instance, SC3 supported 22 clusters. In addition, the large gap between MRtree and initial Seurat clusterings indicated the inability of Seurat to identify accurate and stable clusters on this dataset. This observation was further supported by the lower accuracy (AMRI< 0.6) of the resulting Seurat clusters (Figure S4b).

#### Human pancreas islet cells

To evaluate performance on cell types that are fairly well separated, we investigated the hierarchical structure identified by MRtree for cells from human pancreatic tissues. We first analyzed single-cell RNA sequencing of 635 cells on islets from Wang et al. ^14^, which come from multiple donors, including children, control adults, and individuals with type 1 or type2 diabetes (T1D, T2D). Among them, 430 cells were annotated by the authors into seven cell types, while 205 cells were considered ambiguous and unlabeled. We applied MRtree to construct the hierarchical cluster tree based on SC3 flat clustering with the number of clusters ranging from 2 to 15. The tree was then trimmed to eight leaf nodes based on stability analysis (Figure 3A, Figure S7A). The first split created two large interpretable cell groups: gene ontology (GO) shows enrichment of exocrine functions such as terms related to “Putrescine catabolic process” (adjusted p-value=2.3*E* − 02) and “Cobalamin metabolic process” (adjusted p-value=5.48*E* − 05) for the left branch, and enrichment of endocrine functions such as “Insulin secretion” (adjusted p-value=3.4*E* − 5) and “Enteroendocrine cell differentiation” (adjusted p-value=2.1*E* − 2) for the right branch. The exocrine group was further divided into acinar (*PRSS1*) and ductal cells (*SPP1*). The right branch further separates a previously undiscovered cluster composed mainly of ambiguous cells and a few previously labeled alpha and mesenchyme cells. This cluster expresses marker genes with significant GO terms such as “Collagen metabolic process” and “regulation of endothelial cell migration”, pointing to endothelial and stellate cells (Table S2) that were not labeled in the original analysis. The remaining endocrine cells were further divided into a group containing *α* cells (*GCG*) and pancreatic polypeptide cells (*PPY*), and another group containing *β* (*INS*) and *δ* cells (*RBP4*) (Table S3).

**Figure 3:**
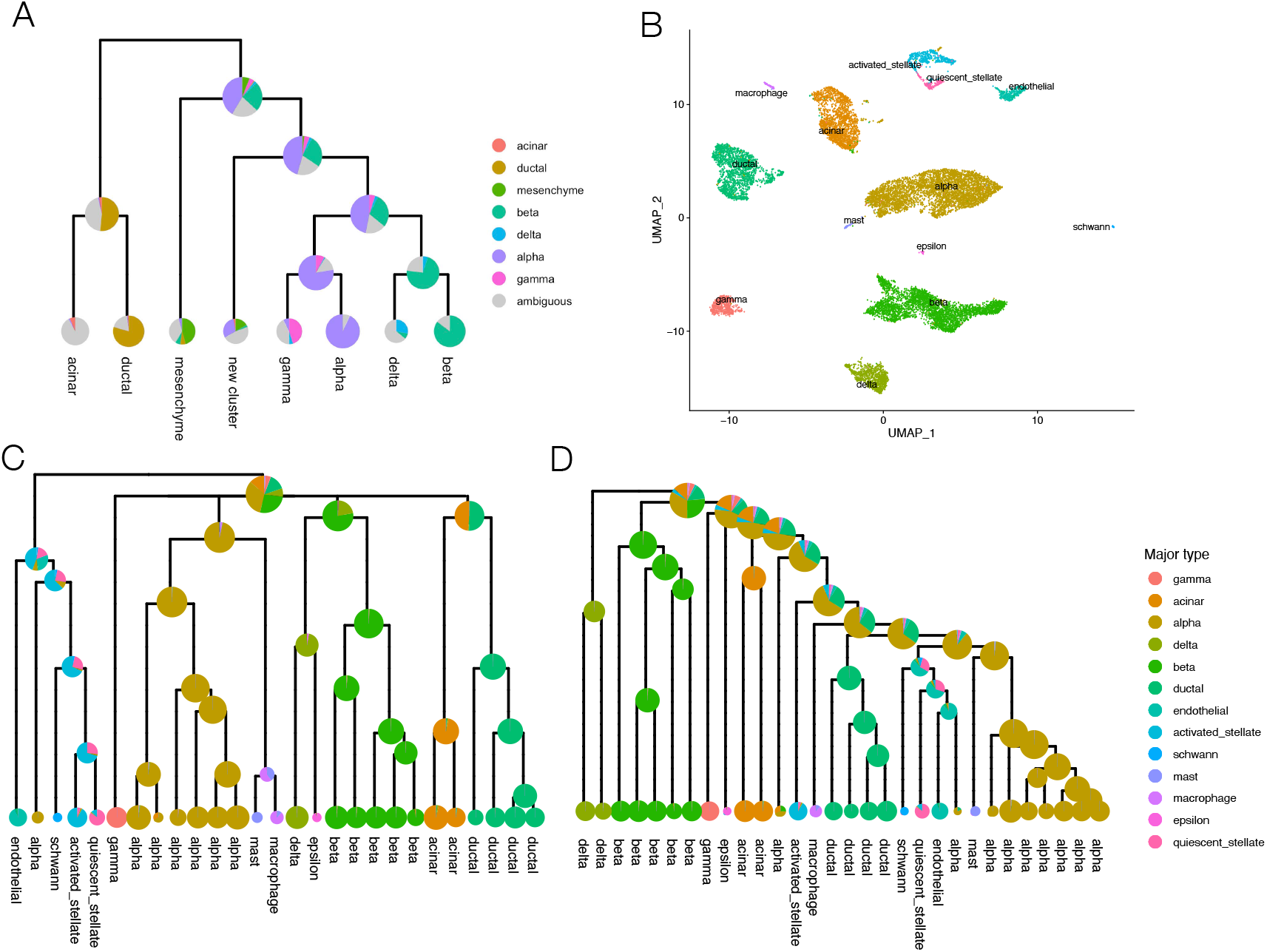
MRtree applied on pancreas islet cells data sets reveals the transcriptional distinctions and similarities between cell types. (A) MRtree-constructed tree with SC3 clusterings on 635 cells from Wang et al. ^14^. The tree was trimmed to the layer with eight leaf clusters (*K* = 8). The pie charts overlying on tree nodes represent the cell type composition for corresponding clusters. Colors indicate the cell-type labels by Wang et al., where a fraction of cells (marked in gray) were considered ambiguous cells by the authors and unlabeled. The leaf labels demonstrate the inferred cluster identity. (B-D) Jointly constructing the cell type hierarchical tree for pancreas islet cells integrated from five technologies. (B) UMAP project of 14,892 cells integrated from five technologies using Seurat MNN integration tools, colored with the cell type labels from respective studies. (C) MRtree-constructed tree from the integrated data with Seurat initial flat clusterings. Pie charts on tree nodes show the cell-type composition given the referencing labels from the studies. Leaf labels indicate the inferred labels of cells in each leaf node. (D) Hierarchical tree constructed by Seurat agglomerative hierarchical clustering starting from Seurat flat clustering results obtained with the highest resolution, annotated similarly by cell type compositions.

In addition to recapitulating a logical tree for all cell types, the eight clusters improved upon the initial SC3 clusters. In particular, seven of the clusters match well with the identified seven major cell types from Wang et al., achieving AMRI greater than 0.95 (Figure S7B-D). By contrast, a competing tree construction method, CellBIC ^6^, revealed a similar tree structure, but it failed to identify the group of *δ* cells ^6^. Finally, because it is well accepted that *β* cells are heterogeneous, especially in conditions of metabolic stress, such as obesity or type 2 diabetes ^14^, we further applied MRtree on the subset of 111 *β* cells. We obtained five *β* subclusters that corresponded to key biological features, including two clusters composed mainly of cells from T2D individuals, and one control group containing 90% cells from children (Figure S7E-G).

Next, we considered a more challenging data set, again from the human pancreatic islet, produced by merging data from five technologies ^8^. In total, 14, 892 cells were annotated and grouped by respective studies into 13 major cell-types with cluster sizes varying by magnitude. We first integrated the cells using Seurat MNN integration tools using 2, 000 highly expressed genes (Figure 3B). Despite the observation that SC3 demonstrates superior performance on the smaller data sets, we utilized Seurat graph-based clustering because it demonstrates greater scalability to large-scale analysis. Flat clusterings were obtained for 50 different resolution parameters sampled via Event-Sampling in the range of [0.001, 2]. The resulting tree identified all 13 major types with high accuracy and also uncovered many subtypes organized as subtrees (Figure 3C). Very distinct cell types separated early and fall into remote branches, while cell types that share similar functions share internal branches and split later in the process. For instance, endothelial, schwann, and stellate cells are very different from other endocrine and exocrine cells and thus split out first. Two types of endocrine cells, acinar and ductal, fall into a common subtree. Likewise, five types of exocrine cells are organized in the same subtree. Finally, subtypes from the same major type are organized in the same subtree, with one exception. A small subset of *α* cells was inappropriately placed in the tree. However, evidence suggests these cells represent an anomaly, possibly due to batch correction. These *α* outliers appear in the UMAP projection separated from other *α* cells and near the activated stellate cells.

For comparison, we produced a hierarchical tree using Seurat agglomerative clustering (Figure 3D). Given the well-separated cluster structure of cell types in the projected PCA space, it is not surprising that the tree also identifies all the major cell types; however, the hierarchical structure appears less reasonable. For instance, the activated and quiescent stellate cells were placed far from each other in the tree, and two endocrine types were grouped in different subtrees. In summary, MRtree produced a more useful tree than competing methods for both applications, and the interpretable subtree structure observed across applications shows promise for further investigation of the cell subtypes identified here.

#### Human fetal brain cells

We applied MRtree to cells from the mid-gestational human cortex, which we call the human brain data^9^. These data were derived from ∼40,000 cells from germinal zones (ventricular zone, subventricular zone) and developing cortex (subplate (SP) and cortical plate (CP)) separated before single-cell isolation. By performing Seurat clustering ^3^, the authors assigned the cells into 16 transcriptionally distinct cell groups (Table S4). For convenience, here we refer to these expert classifications as the Polioudakis labels.

Our analysis began with the same preprocessing steps as conducted in the study ^9^ using the pipeline supported by Seurat V3. The multilevel clustering results are visualized by increasing resolution from the top layer (resolution=0.001) to bottom layer (resolution=2), where each layer corresponds to one clustering (Figure S8A). Notably there were a considerable number of cells assigned to clusters inconsistently over changing resolutions, which made it challenging to determine the optimal resolution and the final cluster memberships. By applying MRtree, we were able to construct the organized hierarchical tree, which was represented by a dendrogram with the cell-type composition of clusters referencing Polioudakis labels shown by pie charts on tree nodes (Figure 4A). MRtree first separated interneurons and pericytes, endothelial and microglia, followed by splitting excitatory deep layer neurons from radial glia to maturing excitatory neurons, representing the rest of a closely connected lineage (upper layer enriched). By further increasing the resolution, the radial glia cells and excitatory neurons were isolated, where the intermediate progenitors were more closely connected with maturing excitatory neurons. The finer distinctions of excitatory neurons were subsequently identified as migrating, maturing, and maturing upper enriched subtypes supported by differential gene expression and canonical cell markers (Table S5). The results were consistent with the group-wise separability visible through a 2-dimensional tSNE projection (Figure 4B). The cluster stability was inspected by comparing the initial Seurat clusters at each resolution with the MRtree results (Figure S8B), which suggested that the clusters were stable up to around *K* = 15. We decided to cut the tree at *K* = 13, which corresponded fairly closely to the 16 major gold standard cell types of the midgestational brain by examination of differentially-expressed marker genes (Table S5, Figure 4C,D). For comparison, an agglomerative hierarchical tree was generated starting from the Seurat clusters obtained at the highest resolution (Figure S8C). These results were distance-based and consequently more vulnerable to outliers, which appear to have caused several anomalies: subsets of ExM, ExM-U, and ExDp1 were grouped together, and two subsets of IP were separated from each other.

**Figure 4:**
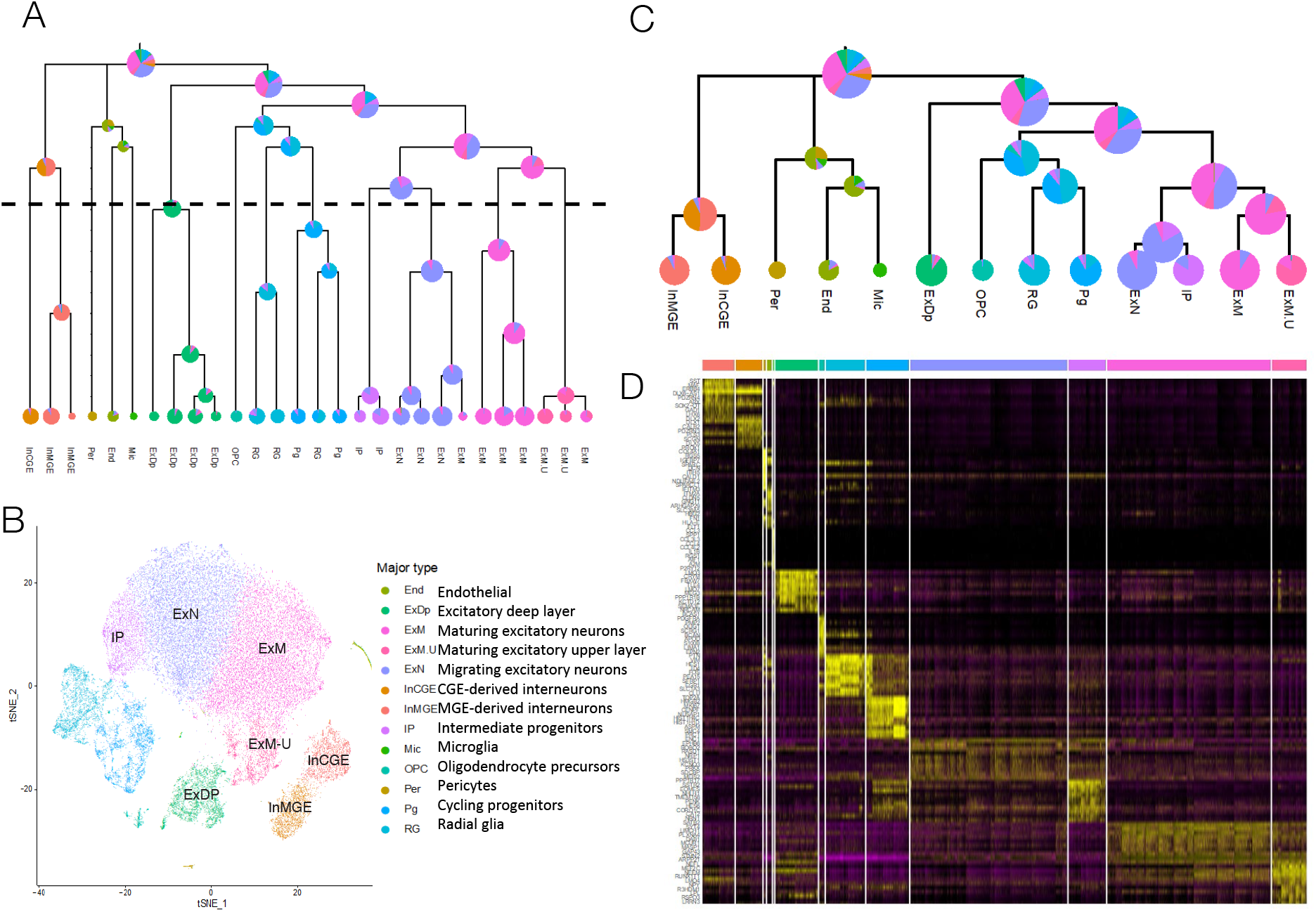
MRtree applied to scRNA-seq data from human brain cells. (A) MRtree produces the hierarchical cluster tree from the initial flat clusterings at multiple resolutions obtained from Seurat V3. The nodes correspond to clusters, with a pie chart displaying the cluster composition referencing the Polioudakis labels. Tree cut is placed at tree layer corresponding to *K* = 13 based on stability analysis, above which the clusters are stable. (B) tSNE plot of all 40,000 human brain cells colored by which of the 13 major clusters the cells belong to, with cell-type identities names hovering over clusters in black. (C) The 13 major clusters were obtained by cutting the tree at the level indicated by the dashed line in (B), indicating the identified major cell types and the associated stable hierarchical structure. (D) Heatmap of the top 10 significant marker genes (FDR-adjusted p-value< 0.05) for the identified 13 major clusters ranked by average log fold change, arranged according to the display of tree leaf nodes.

#### Identify subtypes

Next, we scrutinized the fine-grained structure by re-clustering the 13 major cell types obtained from the hierarchical cluster tree of all cells. The cells were pre-processed from the raw count data as performed in the first iteration, followed by clustering using the Seurat graph-based method. By setting the resolution parameters from 0.05 to 1 and applying MRtree, we obtained one hierarchical tree for each major cell type, determined by trimming the full tree to the stable top layers (ExDP and InMGE are depicted in Figure 5A,B). This resulted in 21 transcriptionally distinct cell types from 7 of the identified major types, expanding IP, ExN, ExM, ExM-U, ExDp, InCGE, and InMGE (Figure 5C, Table S8). The subtypes’ partitions were first evaluated by assessing whether the likely technical and biological co-variation, including brain sample, sequencing run, and cortical region, illustrated somewhat even distribution and appropriate overlap within each identified cluster. Results show that the clusterings were not driven by these technical features and are likely biologically meaningful (Figure 5D, Figure S9).

**Figure 5:**
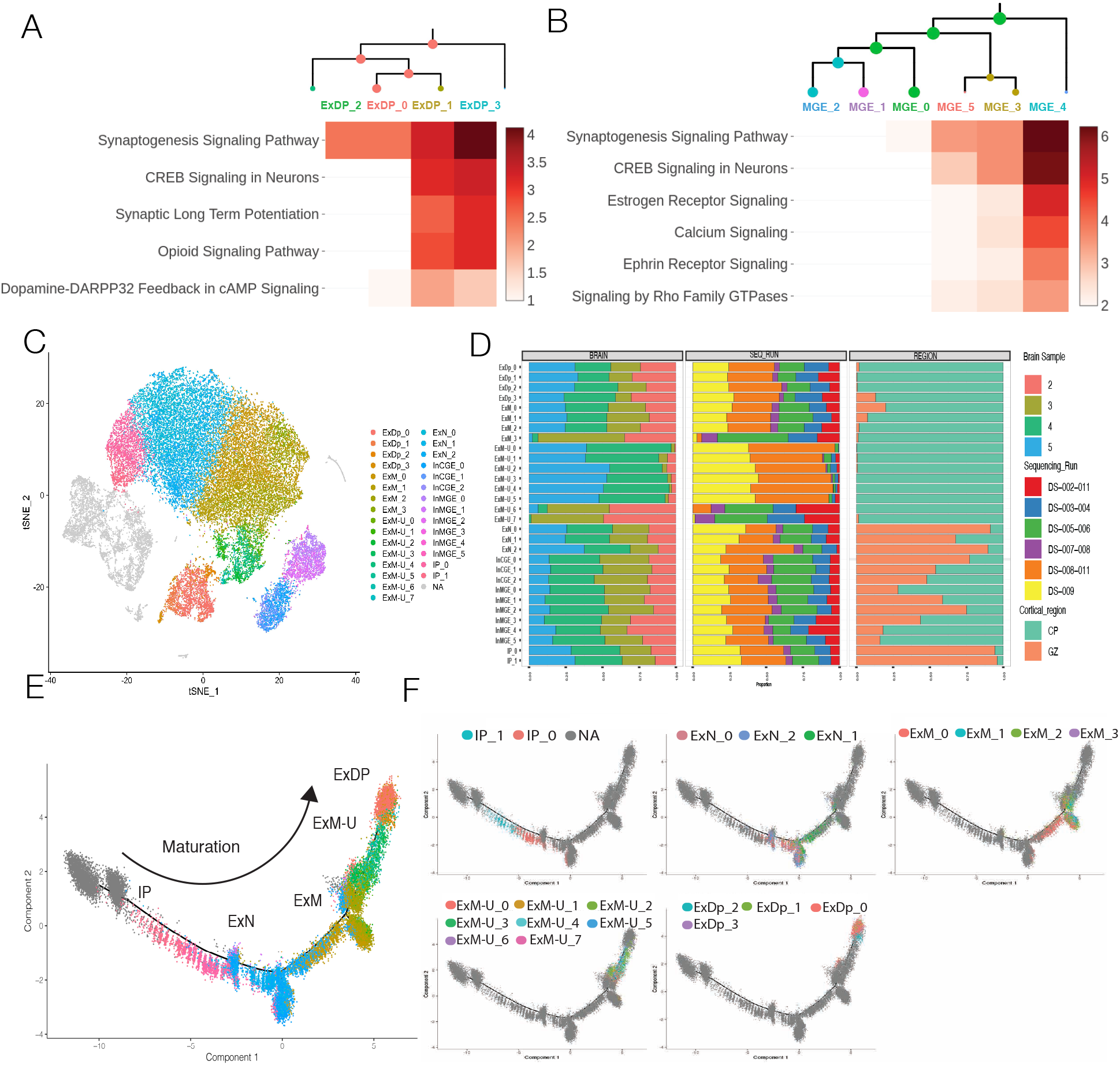
MRtree clusters cells into known cell subtypes and states that underlie known cellular developmental transcriptional trajectories at a higher resolution. (A) Hierarchical cluster tree of subplate/ deep layer excitatory neurons (ExDp) with a heatmap of gene expression within canonical gene ontology categories showing a gradually increased enrichment of Synaptogenesis, CREB signaling, synaptic signaling (i.e., Synaptic long term potentiation, Opioid, and Dopamine-DARPP32-cAMP signaling) across maturation from ExDP 0 to most mature ExDP 3 cluster. (B) Hierarchical cluster tree of MGE-derived interneuron with a heatmap of gene expression within canonical gene ontology categories shows a gradually increased enrichment of Synaptogenesis, CREB signaling, and calcium-mediated signaling across maturation from InMGE 2 to most mature InMGE 4 cluster. (C) tSNE projection of all cells colored by the MRtree identified subtypes from subsequent analysis of the MRtree major cell types. (D) MRtree clusters are driven by biology and not technical co-variation in the data: Histogram of the percentage of cells that each brain sample (left), sequencing run (middle), and cortical region (right) contribute to each cellular cluster identified by MRtree. (E) Cells projected onto Monocle pseudotime analysis from Polioudakis et al., with cells colored by MRtree cell-types and names hovering above. (F) Pseudotime projection of each cluster cell types from MRtree illustrating a continuous developmental trajectory of excitatory neurons, first: top left; intermediate progenitors IP 1, IP 0, top middle; newly born excitatory neurons ExN 0, ExN 2, ExN 1, and, top right; maturing excitatory neurons ExM 0, ExM 1, ExM 2, ExM 3, bottom left; followed by maturing upper layer neurons ExM-U 0 through ExM-U 7 and, bottom left; maturing subplate/ deep layer neurons ExDP 2, ExDP 0, ExDP 1, ExDP 3.

We focus on the results of excitatory neuronal subtypes, given their critical roles in neurological disorders. Close examination revealed that MRtree clustered cells into well-known cell types and states that underlie known cellular developmental transcriptional trajectories at a higher resolution. Projection of each cluster of cell types from MRtree onto Polioudakis Monocle Psuedotime illustrated a continuous developmental trajectory of excitatory neurons, starting from intermediate progenitors (IP) with IP 1 preceding IP 0. The cells then develop into newly born excitatory neurons in the order of ExN 0, ExN 2, ExN 1, which then grow into maturing excitatory neuron subtypes following the order of ExM 0, ExM 1, ExM 2, ExM 3. The trajectory finally ends at maturing upper layer neurons ExM-U 0 through ExM-U 7 and maturing subplate/ deep layer neurons ExDP 2, ExDP 0, ExDP 1, ExDP 3, with ExDP 3 considered as the most mature subtype (Figure 5E,F). The estimated hierarchical tree for subtypes corresponded with gene ontology analysis of differential gene expression between branch cell types. For ExDp, the most distinct subtype was ExDp 3, which was first differentiated from the other subtypes, followed by the split for ExDp 2, and then ExDp 0 and ExDp 1 (Figure 5A). The heatmap of gene expression within canonical gene ontology categories showed a gradual increase in enrichment of Synaptogenesis, *CREB* signaling, synaptic signaling (i.e., Synaptic long-term potentiation, Opioid, and Dopamine-DARPP32-cAMP signaling) across maturation from ExDP 0 to the most mature ExDP 3 cluster. We observed similar functional specializations of inhibitory neuron subtypes (Figure 5B). The most mature subtype InMGE 4 was discriminated from the other MGE interneurons first, followed by splitting the second and third most mature subtypes from less mature cells, and finer distinctions were established subsequently in two branches. The heatmap of gene expression within canonical gene ontology categories showed a gradual increase in enrichment of Synaptogenesis, *CREB* signaling, Calcium mediated signaling across maturation from InMGE 2 to most mature InMGE 4 cluster.

MRtree partitioned intermediate progenitor cells into two subtypes (IP 1 and IP 0; Figure 6A) similar to cell types revealed in Polioudakis et al., achieved only after multiple rounds of analysis of flat clustering results. Marker genes for newly born neurons (i.e. *SLA, STMN2, NEUROD6*) and intermediate progenitors (i.e. *EOMES, SOX11, SOX4, PTN*) showed increased expression markers within IP 0 in contrast to expression of more intermediate progenitors and radial glia genes within IP 1 (i.e. *SLC1A3, VIM, SOX2, HES1*) (Figure 6B). Notably, by comparing the significant protein-protein interacting (PPI) networks from differential genes (DGE) expressed in IP 1 versus significant PPI network from DGE in IP 0, we observed that IP 1 cells PPI contains a highly connected radial glial genes surrounding *VIM* including *MKi67, SOX2*, for example, whereas, IP 0 cells contain more neuronal-committed genes involving early step in neuronal differentiation including *MAPT, GAP43, CALM2, GRIA2, PTPRD* (Figure 6C). Gene ontology analysis further uncovered a switch in the enrichment of *EIF2* signaling, growth factors, and cell cycling pathways (i.e. Sirtuin signaling pathway and *SAPK/JNK* signaling) in IP 1 to more specific neuronal categories like Synaptogenesis, Ephrin Receptor signaling, Reelin signaling underlying migration and neurite pathfinding signaling within IP 0 (Figure 6D).

**Figure 6:**
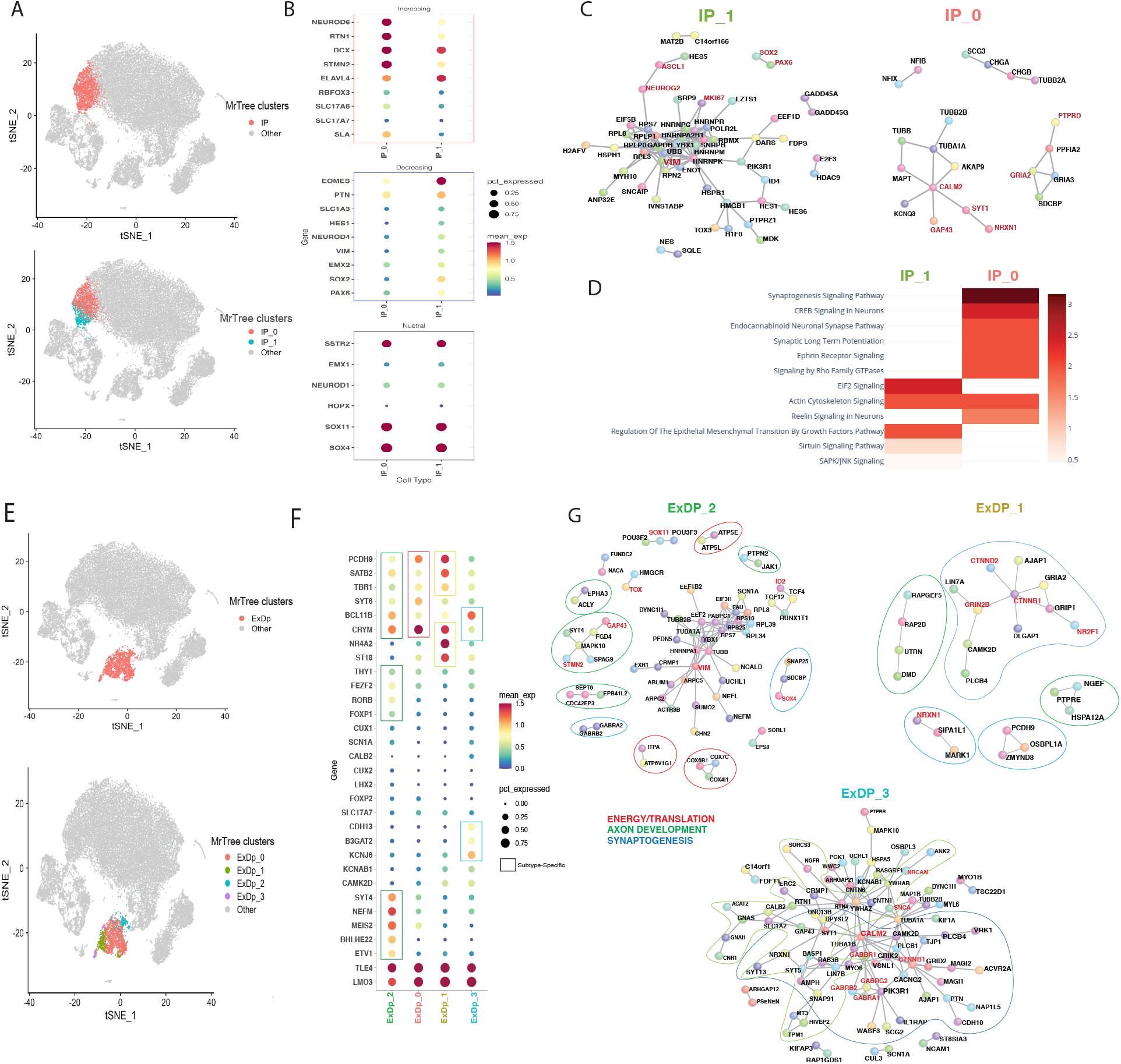
Known and unique biological states identified by MRtree with sub clustering on human fetal brain data: intermediate progenitors and subplate/ deep excitatory neurons (A) top: tSNE plot of all cells where intermediate progenitor (IP) cells identified by MRtree are colored by red, bottom: tSNE projection of MRtree clustering where IP is broken into IP 1, colored in blue and IP 0, colored by red. (B) Gene expression dot plot showing the normalized mean expression of marker genes for newly born neurons (i.e. *SLA, STMN2, NEUROD6*), intermediate progenitors (i.e. *EOMES, SOX11, SOX4, PTN*), and radial glia (i.e. *SLC1A3, VIM, SOX2, HES1*) within IP 1 (left) and IP 0 (right), grouped by increasing (top), decreasing (middle) and neural (bottom) expressions from IP 1 to IP 0. (C) Significant protein-protein interacting (PPI) networks from differential genes expressed in IP 1 on the left versus significant PPI network from DGE in IP 0 on the right. (D) Heatmap of IP 1 and IP 0 gene expression within canonical gene ontology categories. (E) top: tSNE projection where subplate and deep excitatory neurons (ExDP) cells identified by MRtree are colored by red; bottom: tSNE projection where ExDP are broken into ExDP 2, colored in blue and ExDP 0, colored by red, ExDP 1 colored by green, and ExDP 3 colored by purple. (F) Gene expression dot plot showing the normalized mean expression of marker genes for layer 5 (i.e. *ETV1, RORB, FOXP1, FEZF2*), Layer 6 (i.e. *TBR1, SYT6, FOXP2*), shared deep markers (i.e. *RORB, TLE4, LMO3, CRYM, THY1*) and subplate makers (i.e *NR4A2, ST18*) within ExDP 2, ExDP 0, ExDP 1, and ExDP 3 from left to right. The subtype-specific expressions are marked by brackets. (G) Significant protein-protein interacting (PPI) networks from differential genes expressed in ExDP 2 on the top left versus significant PPI network from ExDP 1 top right and PPI from ExDP 3 bottom center.

For ExDP subtypes, a closer examination of the expression of marker genes for layer 5 (i.e., *ETV1, RORB, FOXP1, FEZF2*), Layer 6 (i.e., *TBR1, SYT6, FOXP2*), shared deep markers (i.e., *RORB, TLE4, LMO3, CRYM, THY1*) and subplate makers (i.e., *NR4A2, ST18*) showed expression of Layer 5 markers within the least mature cells, ExDP 2, and layer 6 markers within ExDP 0, in contrast to the more mature expression of layer 6-CTIP2 markers within ExDP 3 and mature expression of markers of subplate and layer 6 within ExDP 1 (Figure 6F). Surprisingly, ExDP 2 PPI revealed a set of genes and structure similar to an intermediate progenitor with *VIM* at the center of translational control and the expression of neuronally committed genes *SOX4, SOX11, ID2* similar to IP 1, except that neuronal specificity genes within this cluster were linked directly to upper Layer 5 cell fate (i.e., *FEZF2, FOXP1, RORB, SYT4*) instead of a general excitatory neuronal lineage seen in IP 1. ExDP 1 subplate cells PPI exhibited a group of connected genes related to more mature cellular properties such as synaptic plasticity and Wnt signaling (i.e., *GRIN2B, CTNNB1, NR2F1, NRXN1*) but no energy or translational pathways that were present in ExDP 2. ExDP 3 cells showed the most extensive and unique PPI that illustrated more committed axonal and synaptic pathways underlying specifically Layer 6 *CTIP2+* cells (i.e., *CALM2, NRCAM, SNCA*, GABAergic postsynaptic machinery) (Figure 6G).

Four other cell types revealed subtypes that were also related to developmental ordering. ExN was partitioned into 3 subtypes that indicate a gradually increased expression of markers of upper layer excitatory neurons in contrast to no expression of deep layer neuronal programs (Figure S10). ExM was partitioned into 4 subtypes, 3 of which illustrate gradually increased expression of upper layer markers, in contrast, a fourth that expressed deep layer markers indicating layer 4/5 excitatory neurons (Figure S11). InMGE was partitioned into 6 progressively more mature subtypes (Fig 5B) that demonstrate distinctions in both maturation and terminal specification (Fig S12). Finally, the 3 subtypes of InCGE display a general maturation of CGE interneurons through a gradual decrease in expression of transcription factors along with a gradual increase in expression of axonal-related genes (Figure S13). Meanwhile, although the signal was sparse, the PPI network for the allegedly most mature subtype revealed a connection between genes critically involved in post-synaptic glutamate signaling and plasticity, further supporting this conjecture. Additional characteristics of these subtypes can be found in Supplemental Information.

## Discussion and conclusion

In this article, we propose MRtree, a computational approach for characterizing multi-resolution cell clusters ranging from major cell groupings to fine-level subtypes using a hierarchical tree. The approach is based on deriving a multi-resolution reconciled tree to integrate clusterings obtained for a range of different resolutions. The proposed method combines the flat and hierarchical clustering results in a novel manner, inheriting the computational efficiency and scalability from the flat clustering and the interpretability of a hierarchical structure. In comparison, MRtree outperforms bottom-up and top-down hierarchical clustering approaches and provides superior clustering for each level of resolution. MRtree also provides tools for sampling implicit resolution parameters for Louvain clustering. This enables equal coverage of different clustering scales as input for the tree construction process. All clustering methods face the challenge of determining the optimal number of clusters supported by the data. While this problem is inherently intractable, MRtree uses a stability criterion to determine the maximum resolution level for which stable clustering results can be obtained for a given dataset. Because MRtree is agnostic to the clustering approach, it can readily utilize input from any flat clustering algorithm. Hence MRtree is extremely flexible, immediately incorporating the advantages of available clustering algorithms, while often providing improved clustering at every resolution due to the reconciliation procedure.

To illustrate the performance of our method, we apply MRtree to a variety of scRNA-seq data sets, including cells from the mouse brain, human pancreas, and human fetal brain tissues. Coupled with suitable initial flat clustering algorithms, MRtree constructs the hierarchical tree that reveals different levels of transcriptional distinction between cell types and outperforms popular competitors, including bottom-up HAC and divisive methods such as CellBIC ^6^. For functionally distinct cell types that can be easily identified, the reconciliation process organizes the clusters obtained under different scales into a unified hierarchical structure, and suggests a proper tree cut to retain the stable partitions. For instance, the constructed tree from integrated pancreatic islet data sets successfully identified endocrine and exocrine groups and subsequent cell types within each group. The clusters from the tree of mouse cortex data sets accurately recovered the known major cell types organized into subtrees of neurons and glial cells. Application of MRtree on human fetal brain cells uncovered previously recognized main types organized in a tree structure along the maturation trajectories.

Our method has a greater impact in challenging situations where clusters are similar. Apart from validating the method on the widely-acknowledged main cell types, we uncovered a list of stable subtypes from the fetal brain dataset that exhibit distinct states and functionality by examining canonical gene ontology categories and significant PPI networks. Specifically, we have shown that two subtypes of intermediate progenitors are well-defined by the expression of radial glia markers versus newly born neurons markers. The subplate /deep layer excitatory neurons are mainly differentiated by the layers the cells will populate. While migrating and maturing excitatory subtypes show a gradual increase of upper layer excitatory neuron markers, upper and deep layer excitatory neuron markers, respectively. Subtypes close in maturation states are reflected in the hierarchical tree as they are split later down the tree. InMGE demonstrates the distinction in both maturation and terminal specification with respect to the engagement of synaptic programs. At the same time, InCGE subtypes differentiate mainly by maturation, which fits nicely with the fact that CGE interneurons are born after MGE interneurons. While both cell types are born in the ventral telencephalon, their terminal specification happens only upon beginning synaptogenesis when they begin to express subtype-specific markers. Surprisingly, subtypes of ExDP revealed a set of genes and structures similar to an intermediate progenitor that can be further investigated in future work. It is worth noting that the quality of MRtree’s construction relies on the performance of the chosen flat clusterings. If the flat clusterings method inputs unstable or biased clusters, these errors will be largely retained and reflected in the estimated hierarchical cluster tree. Similar to many consensus clustering methods, MRtree can be extended to allow input from multiple sources, each applying different flat clustering methods; however, the quality of the constructed tree depends on the clustering performance of the full spectrum of sources. If the input data provides a disparate signal, then the outcome is likely to be unstable.

Our studies suggest several interesting questions worthy of future investigations. For instance, our method is a general framework that allows for any flat-clustering base procedure. In practice, how to determine which base procedure suits better for different data sets still remains open. In addition, the current framework relies on a rough idea about the range of resolution. Can we automatically decide the range of resolution? How can we select this range when the resolution is not parameterized by the number of clusters? In particular, our current method adopts a stability measure to decide whether to further branch the hierarchical tree. Can we provide theoretical guarantees for the power of this stopping criterion? Furthermore, our work shed light on how major cell types evolve to subtypes, and we would like to further verify these biological findings.

## Methods

We briefly formulate the optimization problem and introduce the algorithm we employed. A more detailed description can be found in Supplemental Information. Given the transcriptomes of *n* cells on *p* genes, denoted as *X* ∈ ℝ^*n*×*p*^, suppose clustering is performed using algorithm 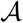 at a range of *m* resolutions with parameters {*k*_1_, …, *k*_*m*_}. Here the resolution parameters are loosely defined where it corresponds explicitly to the number of clusters for some algorithms, while it implicitly determines the number of clusters for other algorithms. To formally state the problem and the hierarchical reconciliation algorithm, we first introduce some notation.

### Definition 1

*A cluster tree T*_*c*_(*k*_1_, …, *k*_*m*_) *at resolution levels* (*k*_1_, …, *k*_*m*_) *is a directed m-partite graph with vertex set V* (*T*_*c*_) *and edge set E*(*T*_*c*_). *Denote the set of all cluster trees as* 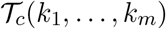.

Here the vertex set *V* (*T*_*c*_) is the union of *m* subsets, namely *V* (*T*_*c*_) = ∪_*j*=1,…,*m*_*V*_*j*_(*T*_*c*_), where each set *V*_*j*_(*T*_*c*_) consists of *k*_*j*_ nodes denoted as {*v*_*j*,1_, …, *v*_*j*_,*k*_*j*_}. Each nodes represents a cluster in the partition of *n* cells into *k*_*j*_ clusters, namely, *v*_*j,k*_ represents the *k*-th cluster at the *j* resolution level. Each direct edge 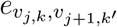 is defined between adjacent layers pointing from a lower resolution cluster *v*_*j,k*_ to a higher resolution cluster *v*_*j*+1,*k′*_ whenever there are overlapping samples between these two clusters at different resolutions. Further, Let *v*_*in*_(*e*) and *v*_*out*_(*e*) be the in-vertex and out-vertex of edge *e*.

### Definition 2

*We call a cluster tree a hierarchical cluster tree, denoted as T*_*h*_(*k*_1_, …, *k*_*m*_), if it satisfies the following constraint:

*Constraint A*_1_: *Each node v*_*j*+1,*k*_ has one and only one in-vertex edge.

*Denote the set of all hierarchical cluster trees at resolution* (*k*_1_, …, *k*_*m*_) *as* 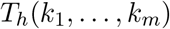

Condition *A*_1_ ensures that any cluster in a higher resolution belongs to one and only one cluster in the adjacent lower resolution, that is, ∀ *v*_*j,k*_, ∃*k′* such that *v*_*j,k*_ ⊆ *v*_*j*−1,*k′*_. It further implies that the hierarchical tree can only be a branching tree as the resolution increases (top-down), and the clusters in lower levels should be intact in levels above it. Compared to the cluster trees, hierarchical cluster trees respect the clustering structure at higher resolutions in the sense that they keep samples that are together at higher resolutions in the same cluster for lower resolutions. Similarly, those samples that are far away from each other at a lower resolution do not enter the same clusters at high resolutions. We illustrate an example of A hierarchical cluster tree and its noisy companion in the form of a cluster tree in Figure S1.

Arrange the the clustering results at each resolution inside a label matrix

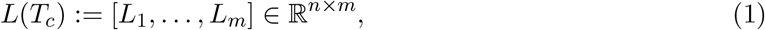

where the *j*-th column denotes the corresponding labels for each data point at resolution *k*_*j*_.

### Definition 3

*For each data point x*_*i*_, *i* = 1, …, *n, define its clustering path p*(*x*_*i*_) := (*v*_1_,*l*_*i1*_, …, *v*_*m*_,*l*_*im*_) *where v*_*j*_,*l*_*ij*_ *is the label for x*_*i*_ *at resolution k*_*j*_. *Let* 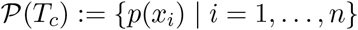 *be the set of all unique paths.*

### Optimization scheme

Let 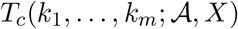 be the initial cluster tree by applying clustering algorithm 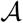 on *X*, and let 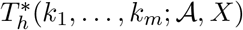 be the underlying true hierarchical cluster tree. Further denote the two respective *n*-by-*p* label matrices as 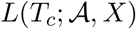 and 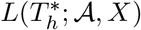. Our goal is to recover the unknown hierarchical tree from the observed initial cluster tree from the multi-resolution flat clustering. For ease of notation, we drop 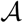, *X* and replace (*k*_1_, …, *k*_*m*_) with **k**^**m**^. Assuming that *T*_*c*_(*k*_*j*_) is an estimator of 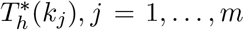, if *T*_*c*_(**k**^**m**^) satisfy constraint *A*_1_, it naturally yields an estimator of 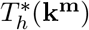, though this is rarely the case. Following this idea, we construct the estimator by building a hierarchical cluster tree that mostly preserves the cluster structures from the observed cluster tree *T*_*c*_(**k**^**m**^) constructed from the initial flat clustering results. To achieve this, we define a loss function as the distance between the solution tree and initial flat clusterings *T*_*c*_(**k**^**m**^). We seek to minimize the loss under the constraint that the solution tree satisfies constraint *A*_1_. To measure the difference between two trees, which is equivalent to measuring mismatch between two sets of partitions, we adopt hamming distance between the respective label matrices (defined in Eq. (1)). Hamming distance computes the number of location-mismatches of a pair of matrices, commonly used for measuring the distance between two paired partitions.

The problem formulated above is equivalent to finding the optimum *k*_*m*_ distinct paths from the set of all feasible paths (Def. 3) of a cluster tree, to which all data points are assigned, and the induced multi-scale partitions preserve the most flat clustering structures. It is a combinatorial optimization problem. The complexity grows exponentially with the depth and number of clusters in each layer of the tree, and therefore is computationally intractable. To alleviate the computational burden, we introduce an equivalent objective function and propose a greedy algorithm to solve it. Formally, define 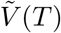 to be the set of “bad” vertices that have more than one in-vertex edge. Then for any proposed hierarchical cluster tree, i.e. 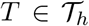, we have 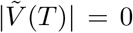. The hierarchy is therefore estimated by solving the optimization problem respective to the newly-formulated constraint,

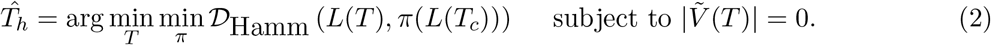

where 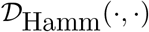 represents the hamming distance. The objective is minimized over permutation of labels 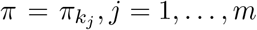 within each partition since the error should not be depending on how we label the classes.

We employ a greedy optimization procedure. The formulated problem (2) is first transformed to a soft constraint problem that shares the same solutions to allow for constraint violation during the optimization procedure. This enables initializing the solution with the observed flat cluster tree *T*_*c*_(**k**_**m**_). The objective is then minimized by sequentially “cleaning” one bad vertex in set 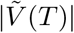 at a time. “Cleaning” the node refers to eliminate all but one edge that have this node as its in-vertex, followed by re-routing data points belonging to the eliminated path to remaining nearest viable paths. The increase in the objective as the results of cleaning the node is considered as the cost of eliminating the node from 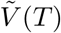. In each iteration, the vertex in set 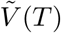 is evaluated for its elimination cost, where the one with the minimum cost is selected. The tree is then updated with the selected node being cleaned and affected data points re-assigned to the nearest remaining paths.

The procedure is repeated until 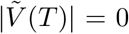, which generates the desired hierarchical cluster tree. The full algorithm is summarized in Algorithm 1. In Supplemental Information we analyze the key properties of the algorithm: Theorem 1 provides the convergence properties, while Theorem 2 describes the memory and time complexity. In addition, we introduce methods for sampling implicit resolution parameters with uniform coverage for modularity-based clustering (Seurat clustering), including linear sampling, exponential sampling, and most preferably, *Event Sampling* method. We also discuss ways of speeding up the algorithm in case of large sample size or a large number of initial flat clusterings through layer-wise reconciliation and performing within-resolution consensus clustering as the first step.

### Stability analysis to determine tree cut

We consider clustering stability to determine the tree cut based on a basic philosophy that clustering should be a structure on the data set that is “stable”. That is, if applied to data sets from the same underlying model, a clustering algorithm should consistently generate similar results. Higher stability across resolutions is reflected as greater consistency of individual initial flat clustering with the resulting clustering in the reconciled tree. To measure the stability, we calculate the similarity using ARI between clusterings in corresponding layers from the initial cluster tree and the resulted hierarchical cluster tree. This will generate a line plot showing the similarity with increasing resolution. The tree cut can then be determined by finding the “change point” where the stability is high at the current point and start to decrease by further increasing the resolution.

### Clustering accuracy

To quantify the clustering performance in each layer of the hierarchical tree, we utilize a novel modified version of Adjusted Rand Index (ARI) ^5^, called Modified Multi-resolution Rand Index (AMRI, Supplemental Information), as the accuracy metric to compare the multi-resolution cluster structures with the true labels. The adjustment allows for comparisons across resolutions, accounting for the reduced ability to uncover details in lower resolutions, thus avoiding a bias towards fine-grained clustering results.

### Tree construction accuracy

To quantify the performance of hierarchical tree construction, given the true tree is known, we reduce the tree to a similarity matrix. Each entry of the matrix represents the length of branch two data points share. The longer branch a pair share, the more similar they are. In this way, we convert the measurement of the difference between hierarchies (dendrograms) to measure the difference between two similarity matrices. The between-similarity distance is measure with the *L*_1_ norm of the difference, defined by

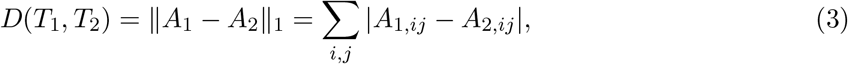

where *A*_1_, *A*_2_ are the similarity matrices of tree *T*_1_, *T*_2_ respectively. Given the certain tree structure, the induced similarity metrics can be visualized in Figure S14.

### Cluster Stability

Apart from examining the performance of MRtree for clustering accuracy, we also access the stability of the clusters at multiple resolutions prior to and post to tree reconciliation. Clustering stability has been considered as a crucial indicator of goodness of the clusters, given that well-performed partitions tend to be consistent across different sampling from the same underlying model or of the same data generating process ^11^. In practice, a large variety of methods has been devised to compute stability scores. Here we adopt the sub-sampling procedure, where the same clustering method is repeatedly performed on the independently sub-sampled data sets and compute the average similarity among the repetitions. Formally given a data set of *n* points *S*_*n*_, let 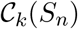 be the resulted clustering outcome with *k* clusters. Let 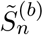 be a sub-sampling of *S*_*n*_ by randomly choosing a subset of size *τn* without replacement. Then the stability score is obtained by averaging the partition similarity on the shared data points,

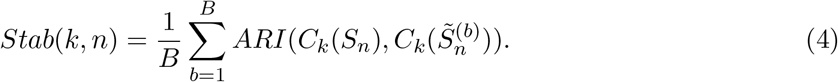

The higher the stability score, the more stable the clustering procedure is regarding the noise in the data. We use *τ* = 0.95 in our experiments.

### Software

MRtree can be constructed using the mrtree R package, which can work directly with Seurat and SingleCellExperiment objects, available on Github (https://github.com/pengminshi/MRtree).

## Declarations

### Ethics approval and consent to participate

Not applicable.

### Consent for publication

Not applicable.

### Availability of data and materials

Not applicable.

### Authors’ contributions

M.P. designed the work, performed the analysis, created the new software used in the work and drafted the work, B.W. performed the analysis and drafted the work, A.E. performed the analysis, D.G. designed the work, Y.W. designed the work and drafted the work, K.R. designed the work and drafted the work.

## Acknowledgements

None at this time.

## References

1. Maayan Baron, Adrian Veres, Samuel L Wolock, Aubrey L Faust, Renaud Gaujoux, Amedeo Vetere, Jennifer Hyoje Ryu, Bridget K Wagner, Shai S Shen-Orr, Allon M Klein, et al. A single-cell transcriptomic map of the human and mouse pancreas reveals inter-and intra-cell population structure. Cell systems, 3(4):346–360, 2016.

2. Florian Buettner, Kedar N Natarajan, F Paolo Casale, Valentina Proserpio, Antonio Scialdone, Fabian J Theis, Sarah A Teichmann, John C Marioni, and Oliver Stegle. Computational analysis of cell-to-cell heterogeneity in single-cell rna-sequencing data reveals hidden subpopulations of cells. Nature biotechnology, 33(2):155, 2015.

3. A. Butler, P. Hoffman, P. Smibert, E. Papalexi, and R. Satija. Integrating single-cell transcriptomic data across different conditions, technologies, and species. Nature Biotechnology, 36(5): 411–+, 2018. ISSN 1087-0156.

4. D. Grun, A. Lyubimova, L. Kester, K. Wiebrands, O. Basak, N. Sasaki, H. Clevers, and A. van Oudenaarden. Single-cell messenger rna sequencing reveals rare intestinal cell types. Nature, 525(7568):251–+, 2015. ISSN 0028-0836.

5. Lawrence Hubert and Phipps Arabie. Comparing partitions. Journal of classification, 2(1): 193–218, 1985. ISSN 0176-4268.

6. Junil Kim, Diana E Stanescu, and Kyoung Jae Won. Cellbic: bimodality-based top-down clustering of single-cell rna sequencing data reveals hierarchical structure of the cell type. Nucleic acids research, 46(21):e124–e124, 2018.

7. V. Y. Kiselev, K. Kirschner, M. T. Schaub, T. Andrews, A. Yiu, T. Chandra, K. N. Natarajan, W. Reik, M. Barahona, A. R. Green, and M. Hemberg. Sc3: consensus clustering of single-cell rna-seq data. Nature Methods, 14(5):483–+, 2017. ISSN 1548-7091.

8. Satija Lab. panc8.SeuratData: Eight Pancreas Datasets Across Five Technologies, 2019. R package version 3.0.2.

9. Damon Polioudakis, Luis de la Torre-Ubieta, Justin Langerman, Andrew G Elkins, Xu Shi, Jason L Stein, Celine K Vuong, Susanne Nichterwitz, Melinda Gevorgian, Carli K Opland, et al. A single-cell transcriptomic atlas of human neocortical development during mid-gestation. Neuron, 103(5):785–801, 2019.

10. Tim Stuart, Andrew Butler, Paul Hoffman, Christoph Hafemeister, Efthymia Papalexi, William M Mauck III, Yuhan Hao, Marlon Stoeckius, Peter Smibert, and Rahul Satija. Comprehensive integration of single-cell data. Cell, 177(7):1888–1902, 2019.

11. Ulrike Von Luxburg. Clustering stability: an overview. 2010.

12. B. Wang, J. J. Zhu, E. Pierson, D. Ramazzotti, and S. Batzoglou. Visualization and analysis of single-cell rna-seq data by kernel-based similarity learning. Nature Methods, 14(4):414–+, 2017. ISSN 1548-7091.

13. X. R. Wang, J. Park, K. Susztak, N. R. Zhang, and M. Y. Li. Bulk tissue cell type deconvolution with multi-subject single-cell expression reference. Nature Communications, 10, 2019. ISSN 2041-1723.

14. Yue J Wang, Jonathan Schug, Kyoung-Jae Won, Chengyang Liu, Ali Naji, Dana Avrahami, Maria L Golson, and Klaus H Kaestner. Single-cell transcriptomics of the human endocrine pancreas. Diabetes, 65(10):3028–3038, 2016.

15. Joe H Ward Jr. Hierarchical grouping to optimize an objective function. Journal of the American statistical association, 58(301):236–244, 1963.

16. Nicola K Wilson, David G Kent, Florian Buettner, Mona Shehata, Iain C Macaulay, Fernando J Calero-Nieto, Manuel Sánchez Castillo, Caroline A Oedekoven, Evangelia Diamanti, Reiner Schulte, et al. Combined single-cell functional and gene expression analysis resolves heterogeneity within stem cell populations. Cell stem cell, 16(6):712–724, 2015.

17. Luke Zappia and Alicia Oshlack. Clustering trees: a visualization for evaluating clusterings at multiple resolutions. GigaScience, 7(7):giy083, 2018.

18. Amit Zeisel, Ana B Muñoz-Manchado, Simone Codeluppi, Peter Lönnerberg, Gioele La Manno, Anna Juréus, Sueli Marques, Hermany Munguba, Liqun He, Christer Betsholtz, et al. Cell types in the mouse cortex and hippocampus revealed by single-cell rna-seq. Science, 347(6226): 1138–1142, 2015.

19. X. W. Zhang, C. L. Xu, and N. Yosef. Simulating multiple faceted variability in single cell rna sequencing. Nature Communications, 10, 2019. ISSN 2041-1723.

20. L. X. Zhu, J. Lei, B. Devlin, and K. Roeder. A unified statistical framework for single cell and bulk rna sequencing data. Annals of Applied Statistics, 12(1):609–632, 2018. ISSN 1932-6157.

21. Lingxue Zhu, Jing Lei, Lambertus Klei, Bernie Devlin, and Kathryn Roeder. Semisoft clustering of single-cell data. Proceedings of the National Academy of Sciences, 116(2):466–471, 2019.

22. J. Zurauskiene and C. Yau. pcareduce: hierarchical clustering of single cell transcriptional profiles. Bmc Bioinformatics, 17, 2016. ISSN 1471-2105.

