## Supplemental Information for "Cell Type Hierarchy Reconstruction via Reconciliation of Multi-resolution Cluster Tree"

**2 Adjusted Multi-resolution Rand Index (AMRI)** **10**

**3 Simulated data** **11**

**4 Single RNA-seq data analysis** **13**

### 1 Algorithm details

Let $T_c(k_1, \ldots, k_m; \mathcal{A}, X)$ be the initial cluster tree by applying clustering algorithm $\mathcal{A}$ on $X$, and let $T_h^*(k_1, \ldots, k_m; \mathcal{A}, X)$ be the underlying true hierarchical cluster tree. Further denote the two respective $n$-by-$p$ label matrices as $L(T_c; \mathcal{A}, X)$ and $L(T_h^*; \mathcal{A}, X)$. Our goal is to recover the unknown hierarchical tree from the observed initial cluster tree from the multi-resolutional flat clustering. For ease of notation, we drop $\mathcal{A}, X$ and replace $(k_1, \ldots, k_m)$ with $\mathbf{k^m}$. A natural estimator finds the hierarchical tree that preserves the most cluster structures from the observed cluster tree $T_c(\mathbf{k^m})$ constructed by the initial flat clustering results. More formally, if we define a distance metric $D(\cdot, \cdot)$ that measures the discrepancy between two cluster trees, the ideal estimator minimizes the distance of $\hat{T}_h(\mathbf{k^m})$ from $T_c(\mathbf{k^m})$, namely

$$\hat{T}_h(\mathbf{k^m}) := \arg \min_{T_h \in \mathcal{T}_h(\mathbf{k^m})} D\big(T_h, T_c(\mathbf{k^m})\big). \tag{5}$$

where $\mathcal{T}_h(\mathbf{k^m})$ denotes the set of hierarchical cluster trees with resolution $\mathbf{k^m} := (k_1, \ldots, k_m)$ (satisfying the constraint $A_1$).

A common way of measuring the distance between matched partitions is to calculate the hamming distance (or pair-counting distance) between the respective label matrices (defined in Eq (1) in main text), namely

$$\begin{aligned} D\big(T_h(\mathbf{k^m}), T_c(\mathbf{k^m})\big) &= \min_{\pi} \mathcal{D}_{\text{Hamm}}\left(L(T_h), \pi(L(T_c))\right) \\ &= \sum_{k=1}^{m} \min_{\pi_k} \mathcal{D}_{\text{Hamm}}\left(l_k(T_h), \pi_k\big(l_k(T_c)\big)\right) \\ &= \sum_{k=1}^{m} \min_{\pi_k} \sum_{i=1}^{n} 1_{\{l_{ik}(T_h) \neq \pi_k(l_{ik}(T_c))\}}, \end{aligned} \tag{6}$$

where $\mathcal{D}_{\text{Hamm}}(\cdot, \cdot)$ represents the hamming distance between two matrices or vectors, which computes the number of location-mismatches. Here the objective is to minimize over all the label permutations $\pi_k$ within layer $k, k = 1, \ldots, m$, because the error should not depend on how we label the classes.

Note that the optimization problem formulated above (5 and 6) is a combinatorial optimization problem, which is often computationally infeasible to solve. This can be easily seen with the following properties for hierarchical cluster trees.

*(i)* A cluster tree contains at most $\prod_{j=1}^{m} k_j$ distinct paths, and a hierarchical cluster tree contains at most $k_m$ distinct path, namely

- $\forall T_c \in \mathcal{T}_c(k_1, \ldots, k_m)$, we have $|\mathcal{P}(T_c)| \leq \prod_{j=1}^{m} k_j$,
- $\forall T_h \in \mathcal{T}_h(k_1, \ldots, k_m)$, we have $|\mathcal{P}(T_h)| \leq k_m$.

*(ii)* In a hierarchical cluster tree $H$, there is no merging path in $\mathcal{P}(T_h)$, i.e. $\forall p_s, p'_s \in \mathcal{P}(T_h)$, if $\exists j$ such that $p_s(j) \neq p_{s'}(j)$, then $p_s(l) \neq p_{s'}(l)$ for all $l \geq j$.

Indeed, for an initial cluster tree with number of clusters increasing from $k_1$ to $k_m$, the set of all feasible path has cardinality $k_m!/(k_1 - 1)!$. Inferring the optimal solution of (5) is equivalent to selecting an optimal set of at most $k_m$ path such that they satisfy the constraints of a hierarchical cluster tree, i.e., a partition tree, while minimizing the objective in (5). Regardless of the label permutation, the computational cost for exhaustive search is $nk_m\binom{k_m!/(k_1-1)!}{k_m}$, which is intractable for a deep tree. To alleviate the computational burden, we propose a greedy algorithm to approximate the solution.

#### 1.1 A greedy algorithm

Instead of searching for an optimal hierarchical tree in an exhaustive manner, in this section, we propose an optimization algorithm that constructs the tree by sequentially expanding the branches and updating the branch assignment of each data point.

First, recall that the optimization problem (5) is defined as minimizing the hamming distance between the label matrix $L(T)$ and $L(T_c)$ subject to the constraint that $T \in \mathcal{T}_h$. If we define $\tilde{V}(T)$ to be the set of "bad" vertices that have more than one in-vertex edges, then for any $T$ being a hierarchical cluster tree, we have $|\tilde{V}(T)| = 0, T \in \mathcal{T}_h$. Therefore the optimization scheme (5) is equivalent to the following problem

$$\min_{T} \min_{\pi} \mathcal{D}_{\text{Hamm}}\left(L(T), \pi(L(T_c))\right) \quad \text{subject to } |\tilde{V}(T)| = 0. \tag{7}$$

Introducing a penalization parameter $\lambda > 0$, we instead consider an associated optimization problem

$$\min_T \min_\pi J(T) := \mathcal{D}_{\text{Hamm}}\left(L(T), \pi(L(T_c))\right) + \lambda \sum_{v \in V(T)} 1_{v \in \tilde{V}(T)}. \tag{8}$$

It is easily seen that the solution to the above problem is the same as that of the original optimization problem (7) when $\lambda > nm$, the penalty term will dominate the first term otherwise.

**Optimization procedure** Now we are ready to describe our method of finding the optimal solution to the above optimization problem. We utilize an iterative procedure that greedily minimizes the objective function (Eq. (8)) at each step. We initialize the solution with the original cluster tree $C$, denoted as $H^{(0)}$ and then proceed sequentially by correcting one "bad" vertex at a time in the steepest descent direction of the objective function. To decide which node to correct next, we calculate the change of the objective function $J(T)$ for correcting each bad vertex by eliminating all but one in-edge of that vertex. Namely, for each edge $e(v_{j-1,l'}, v_{j,l})$, we compute the effect to the objective function if we eliminate all its conflicting edges,

$$\Delta_{e(v_{j-1,l'},v_{j,l})} J(T)\big|_{T^{t-1}} := \mathcal{D}_{\text{Hamm}}\left(L(T_c), L(\tilde{T}^{(t-1)})\right) - \mathcal{D}_{\text{Hamm}}\left(L(T_c), L(T^{(t-1)})\right) - \lambda. \tag{9}$$

Intuitively, finding the edge $e(v_{j-1,l'}, v_{j,l})$ that minimizes $\Delta_{e(v_{j-1,l'},v_{j,l})} J(T)\big|_{H^{t-1}}$ corresponds to finding the steepest descent direction.

In order to understand the above quantity, some notations are in order.

- Quantity $\tilde{T}^{(t-1)}$ is the one-step-edited graph from $T^{(t-1)}$ by removing all the conflicting edges of $e(v_{j-1,l'}, v_{j,l})$, so that

$$\begin{aligned} &E(\tilde{T}^{(t-1)}) = E(T^{(t-1)}) \setminus \cup_{\{e: v_{\text{in}}(e) = v_{i,l'}, e \neq e(v_{j-1,l}, v_{j,l'})\}} e, \\ &V(\tilde{T}^{(t-1)}) = V(T^{(t-1)}). \end{aligned} \tag{10}$$

- Quantity $L(T^{(t-1)})$ is the result of re-assigning data points that are affected by the edge elimination to the remaining paths in $T^{(t-1)}$ given by

$$L(T^{(t-1)}) = \Theta^{(t-1)} P(T^{(t-1)}), \quad \Theta^{(t-1)} = \text{argmin}_{\Theta} \mathcal{D}_{\text{Hamm}} \left( L(T_c), \Theta P(T^{(t-1)}) \right), \tag{11}$$

where $P(T^{(t-1)}) \in \mathbb{R}^{n_p^{(t-1)} \times m}$ is the set of all viable paths in $T^{(t-1)}$ arranged column-wisely and $n_p^{(t)}$ is the total number of distinct viable paths.

- Quantity $\Theta^{(t-1)} \in \mathbb{R}^{n \times n_p^{(t-1)}}$ is a binary membership matrix representing the assignment of node to one of the path, $\Theta_{ij} \in \{0, 1\}$ and $\Theta^{(t-1)} \mathbf{1}_{\mathbf{n_p^{(t-1)}}} = \mathbf{1_n}$. $L(\tilde{T}^{(t-1)})$ is obtained following similar procedure where the number of viable paths becomes $\tilde{n}_p^{(t-1)}$ in $\tilde{T}^{(t-1)}$ (The permutation is omitted since the permutation is intrinsically determined during optimization process).

Note that $\tilde{n}_p^{(t-1)} < n_p^{(t-1)}$ since $E(\tilde{T}^{(t-1)}) \subset E(T^{(t-1)})$. The previous step guarantees that the vertex $v_{j,l}$ now has only one in-edge $e(v_{j-1,l'}, v_{j,l})$, thus can be successfully removed from the bad vertex set while subsequently reducing the objective function by $\lambda$ minus the difference in the hamming distance. Therefore trimming the tree towards its steepest descent direction is then achieved by selecting among in-edges of bad vertices and removing all its conflicting edges. More specifically, if we define the cost associated to each edge at iteration $t$ as

$$w^{(t)}(e(v_{j-1,l}, v_{j,l'})) := \mathcal{D}_{\text{Hamm}} \left( L(T_c), L(\tilde{T}^{(t-1)}) \right) - \mathcal{D}_{\text{Hamm}} \left( L(T_c), L(T^{(t-1)}) \right), \tag{12}$$

then the steepest descent is obtained by minimizing this cost, namely

$$e^{(t)} := \text{argmin}_{e: v_{out}(e) \in \tilde{V}(T^{(t)})} w^{(t)}(e). \tag{13}$$

To summarize, at iteration $k$, the bad vertex set $\tilde{V}(T^{(t-1)}$ is located, and the costs for each in-edge of the bad vertices are calculated, among which the minimum is selected, and all its conflicting edges are removed — giving an updated tree $T^{(t)}$. Then the data points are reassigned to the remaining viable paths that maintain the most similarity compared to initial multi-resolution clustering, producing the new label matrix $L(T^{(t)})$. This process is repeated until $\tilde{V}(T^{(t)}) = \emptyset$. For easy reference, we summarize the entire optimization procedure in algorithm 1.

**Theoretical properties** We are ready to state some properties of the proposed algorithm. Theorem 1 provides the convergence property and while Theorem 2 describes the memory and time complexity.

---

**Algorithm 1:** MRtree

---

**Input:** Initial cluster tree $T_c(\mathbf{k^m})$ (label matrix $L(T_c)$)

**Output:** Reconstructed hierarchical cluster tree $\hat{T}_h(\mathbf{k^m})$ (label matrix $L(\hat{T}_h)$)

`/* Initialization */`

$H^{(0)} = T_c$

$\tilde{V}^{(0)} = \tilde{V}(T^{(0)}),\ P^{(0)} = P(T_c)$

`/* Start iteration */`

t=1

**while** $\tilde{V}^{(t-1)} \neq \emptyset$ **do**

- Calculate the edge costs by (12)
- Select the optimum edge $e^{(t)} = e(v_{j-1,l'}, v_{j,l})$ by (13)
- Eliminate the conflicting edges of $e^{(t)}$ to obtain $T^{(t)}$
- Update $\tilde{V}^{(t)} = \tilde{V}(T^{(t)}), P^{(t)} = P(T^{(t)})$
- Re-assign the data point to the viable path $P^{(t)}$ by (11) to obtain updated label matrix $L(T^{(t)})$.
- $t = t + 1$

$\hat{T}_h(\mathbf{k^m}) = T^{(t)}, L(\hat{T}_h) = L(T^{(t)})$ are the final hierarchical cluster tree and corresponding label matrix of the data points.

---

**Theorem 1** *(Convergence property) Algorithm 1 satisfies the following*

- $|\tilde{V}(H^{(t)})| < |\tilde{V}(H^{(t-1)})|,\ \forall t,$
- $|P^{(t)}| < |P^{(t-1)}|,\ \forall t,$

*which implies that the procedure converges to an local minimum in at most* $|\tilde{V}(H^{(0)})| = \tilde{V}(C)$ *steps.*

**Theorem 2** *(Complexity) Algorithm 1 has time complexity* $O(k_m^{\alpha} nm) \approx O(k_m m^{\alpha+1}), \alpha \leq 3$, *where* $k_m$ *is the number of clusters in the highest resolution,* $m$ *is the number of layers and* $n$ *is number of data points. The worst case* $\alpha = 3$ *is achieved when the initial cluster tree is a* $m$*-partite fully connected graph. The memory requirement is* $O(mn)$.

#### 1.2 Sample resolutions with uniform coverage

To build the hierarchical cluster tree and preserve the features of clusterings at different scales, we want to sample a range of resolution parameters in a way that ensures we give equal coverage to different clustering scales. For methods explicitly requiring $K$, the number of clusters, this can be achieved by conducting an equal number of clusterings for each $K$. For method such as Louvain with an implicit resolution parameter $\gamma$, this becomes a non-trivial task as the possible values for the resolution parameter can often span several orders of magnitude, and the sensitivity of obtained community structure on the value of the resolution parameter varies widely depending on the value of the resolution parameter itself. Two obvious sampling strategies would be linear sampling and exponential sampling. For linear sampling, one would use equally spaced values of $\gamma$ between $\gamma_{\min}$ and $\gamma_{\max}$, and for exponential sampling, one would use values that are equally spaced on a logarithmic scale. In addition to the two methods described above, we also implemented the Event Sampling method [2]. Event Sampling directly exploits the behavior of the modularity quality function to provide good coverage of different scales in a network. Given the adjacency matrix $A$ and the labels vector $\vec{g}$, consider the modularity function

$$Q(\vec{g},\gamma) = \sum_{i,j=1}^{n} \left(A_{ij} - \gamma P_{ij}\right) \delta\left(g_i, g_j\right), \quad P_{ij} = \frac{k_i k_j}{2m}, \quad k_i = \sum_j A_{ij}, \quad 2m = \sum_i k_i. \tag{14}$$

The relative magnitude of the antiferromagnetic interactions [2] was shown to be a good approximation of clustering scales [2],

$$\beta(\gamma) = \frac{\sum_{(i,j)\in E^-(\gamma)} |A_{ij} - \gamma P_{ij}|}{\sum_{(i,j), i\neq j} |A_{ij} - \gamma P_{ij}|}, \quad E^-(\gamma) = \left\{(i,j) \mid i \neq j, A_{ij} - \gamma P_{ij} < 0\right\}, \tag{15}$$

where $\beta(\gamma)$ is monotonically increasing for $\gamma$. Therefore resolution $\gamma$ can be sampled through inverting the relationship between $\gamma$ and $\beta$

$$\gamma(\beta) = \frac{\sum_{(i,j)\in E^-(\beta)} A_{ij} + \beta\left(\sum_{(i,j)\in E^+(\beta)} A_{ij} - \sum_{(i,j)\in E^-(\beta)} A_{ij}\right)}{\sum_{(i,j)\in E^-(\beta)} P_{ij} + \beta\left(\sum_{(i,j)\in E^+(\beta)} P_{ij} - \sum_{(i,j)\in E^-(\beta)} P_{ij}\right)}, \tag{16}$$

and sample $\gamma(\beta)$ at equally spaced values of $\beta$ between $\beta_{\min}$ and $\beta_{\max}$.

The inversed relationship can be solved via identifying the distinct values of $\sum_{(i,j)\in E^{-}(\gamma)} A_{ij}$ with increasing $\gamma$ and the use of interpolation. Due to the high computational complexity, we first randomly subsample a $m$-by-$m$ sub-matrix from $A$, and then perform the Event Sampling for the sub-matrix. We note that the sampled resolutions show little variance when a sub-matrix is used.

We examined the performance of Event Sampling with Louvain clustering using the simulated data from SymSim package. In total 500 cells were generated from 8 types given an assumed tree structure. To make sure the parameters were sampled from the same weighted adjacent matrix used for clustering, we extracted the shared nearest neighbor graph from the Seurat `FindNeighbors` results. 20 resolution parameters were sampled with $\gamma \in (0.01, 3)$. Figure S5 visualizes the number of clusters produced for each resolution versus the order of resolution over repeated simulations. We achieved an approximately linear relationship between the index and the number of clusters, indicating that the resolutions were sampled uniformly across scales.

#### 1.3 Layer-wise algorithm speed-up and within-resolution consensus clustering

**Top-down layer-wise reconciliation** The complexity of the reconciliation algorithm 1 scales polynomially with the number of layers in the cluster tree, i.e., the total number of clustering. In cases of a large number of layers, a top-down layer-wise reconciliation can be employed instead to speed up the process, where in each iteration, the selection of node with minimum cost is constrained to the top few layers. It is observed in empirical experiments that select candidate node from the top two layers is sufficient to generate good results.

**Within-resolution consensus clustering** It is worth noticing that this layer-wise reconciliation process is more susceptible to suboptimum if some of the clusterings deviate far away from the true partition. The problem can be alleviated with an initial consensus clustering step if multiple clusterings exist with the same $K$, which further speed up the reconciliation process. We call this procedure *within-resolution consensus clustering.* This procedure can be employed when the clustering involves stochasticity for the fixed $K$, for instance, when multiple different resolution parameters are mapped to the same $K$, as the case for Louvain method, and when using K-means for an intermediate step for clustering that can generate local optimum and thus rely on random seed chosen as the case of SC3[4] and SOUP[14].

Compared to across resolution reconciliation, within-resolution consensus clustering is an easier problem, especially when the number of clusters in each partition is the same and equal to the final number of clusters in the consensus partition[10]. Many methods have been proposed utilizing different consensus functions. Here we adopt a fast method described as follows.

The input of the consensus procedure is a collection of $m_k$ clusterings each composed of $k$ clusters with labels $l_j^{(k)} \in [k]^n, j = 1, \dots, m_k$. We aim to seek a consensus clustering with the labels $l^{(k)}$ that minimize the Hamming distance

$$l^{(k)} = \arg \min_{l \in [K]^n} \min_{\sigma = \left(\sigma_1, \dots, \sigma_{m_k}\right)} \sum_{j=1}^{m_k} \mathcal{D}_{\text{Hamm}} \left(\sigma_j(l_j^{(k)}), l\right) \tag{17}$$

where $\sigma_j$ are bijections from $[k]$ to $[k]$. Denote the respective binary membership matrix as $H_j^{(k)}, j = 1, \dots, m_k$, such that $H_j^{(k)}(i, s) = 1$ if $l_j^{(k)}(i) = s$, otherwise 0. Then we have the equivalent form of

$$\begin{aligned} H^{(k)} &= \arg \min_{H \in \mathcal{M}(n,k)} \min_{\Gamma_1, \dots, \Gamma_{m_k}} \sum_{j=1}^{m_k} \left\| H_j^{(k)} - H\Gamma_j \right\|_F^2 \\ &= \arg \min_{H \in \mathcal{M}(n,K)} \min_{\Gamma = \left[\Gamma_1, \dots, \Gamma_{m_k}\right]} \left\| [H_1^{(k)}, \dots, H_{m_k}^{(k)}] - H\Gamma \right\|_F^2 \end{aligned} \tag{18}$$

where $\Gamma_j, j = 1, \dots, m_k$ are column permutation matrices, and $\mathcal{M}(n, k)$ represents the set of all $n$-by-$k$ binary membership matrix. To recover $H^{(k)}$, we perform truncated SVD on $\left[H_1^{(k)}, \dots, H_{s_k}^{(k)}\right]$ to obtain the top-$k$ left singular vectors. The singular vectors are weighted by corresponding singular values, followed by a K-means clustering. The resulting $k$ clusters give the consensus clustering result. This procedure is computation and memory-efficient since it does not require constructing the $n$-by-$n$ co-classification matrix. When $k \ll n$, fast algorithms can achieve time complexity of roughly $O\left(T(nm_k k + k^3 + c)\right)$, where $T$ is the number of iterations, and $c$ is the other computational cost for each step.

#### 2 Adjusted Multi-resolution Rand Index (AMRI)

In this section, we describe a novel criterion to evaluate the difference between clustering results obtained with different resolutions, which is referred to as the Adjusted Multi-resolution Rand Index (AMRI). The criterion is based on the prior works[7,1] that measures the similarity between flat clustering results and is adjusted to account for the difference in clustering resolutions.

Given two sets of clustering results $\mathcal{A} = \{A_1, \ldots, A_{K_\mathcal{A}}\}, \mathcal{B} = \{B_1, \ldots, B_{K_\mathcal{B}}\}$ of data set $S$ that consists $n$ elements, the element pairs can be classified into four subsets:

$$\begin{aligned}
S_{\mathcal{A},\mathcal{B}} &= \{\text{pairs in } S \text{ that are in the same cluster in } \mathcal{A} \text{ and in the same cluster in } \mathcal{B}\},\\
S_{\mathcal{A}^c,\mathcal{B}^c} &= \{\text{pairs in } S \text{ that are in different clusters in } \mathcal{A} \text{ and in different clusters in } \mathcal{B}\},\\
S_{\mathcal{A},\mathcal{B}^c} &= \{\text{pairs in } S \text{ that are in the same cluster in } \mathcal{A} \text{ but in different clusters in } \mathcal{B}\},\\
S_{\mathcal{A}^c,\mathcal{B}} &= \{\text{pairs in } S \text{ that are in different clusters in } \mathcal{A} \text{ but in the same cluster in } \mathcal{B}\}.
\end{aligned}$$

The Rand Index (RI)[7] is a well-known criterion to measure the similarity between the two clustering results, which is defined as

$$RI(\mathcal{A},\mathcal{B}) := \frac{|S_{\mathcal{A},\mathcal{B}}| + |S_{\mathcal{A}^c,\mathcal{B}^c}|}{|S_{\mathcal{A},\mathcal{B}}| + |S_{\mathcal{A}^c,\mathcal{B}^c}| + |S_{\mathcal{A},\mathcal{B}^c}| + |S_{\mathcal{A}^c,\mathcal{B}}|} = \frac{|S_{\mathcal{A},\mathcal{B}}| + |S_{\mathcal{A}^c,\mathcal{B}^c}|}{\binom{n}{2}}. \tag{19}$$

It is easily seen that the quantity $RI(\mathcal{A},\mathcal{B})$ lies between 0 and 1, where 1 indicates the two clusterings are identical and 0 occurs for clusters which do not share a single pair of elements. The RI is then adjusted by chance to produce Adjusted-RI (ARI)[1]. By far, the most common approach to correct clustering similarity for chance assumes that both clusterings are uniformly and independently sampled from the permutation model, so that

$$\begin{aligned}
ARI_{\text{perm}}(\mathcal{A},\mathcal{B}) &= \frac{RI(\mathcal{A},\mathcal{B}) - \mathbb{E}_{\text{perm}}[RI(\mathcal{A},\mathcal{B})]}{\max(RI) - \mathbb{E}_{\text{perm}}[RI(\mathcal{A},\mathcal{B})]} = \frac{RI(\mathcal{A},\mathcal{B}) - \mathbb{E}_{\text{perm}}[RI(\mathcal{A},\mathcal{B})]}{1 - \mathbb{E}_{\text{perm}}[RI(\mathcal{A},\mathcal{B})]},\\
\mathbb{E}_{\text{perm}}[RI(\mathcal{A},\mathcal{B})] &= \frac{\sum_i \binom{|A_i|}{2}}{\binom{n}{2}} \frac{\sum_j \binom{|B_j|}{2}}{\binom{n}{2}} + \Big(1 - \frac{\sum_i \binom{|A_i|}{2}}{\binom{n}{2}}\Big)\Big(1 - \frac{\sum_j \binom{|B_j|}{2}}{\binom{n}{2}}\Big).
\end{aligned} \tag{20}$$

The formula is equivalent for both one-sided and two-sided model.

To see why we need a modified accuracy metric, consider a toy example where there are $K$ underlying clusters. Suppose there are $K-1$ underlying clusters and a clustering algorithm successfully identify the first $K-2$ clusters and group the other two underlying clusters together, the ARI of the produced partition is large due to the failure of combing two underlying clusters. However, one may fix the problem by re-clustering the last group with $K=2$ to recover the other labels. This example suggests that the original ARI may not be suitable if one cares about more than one shot of the clustering such that the clustering results are obtained at different resolutions.

Taking this resolution factor into account, we propose the Adjusted Multi-resolution Rand Index (AMRI), adopting the same notion as that of ARI. Without loss of generality, assume that $K_{\mathcal{A}} < K_{\mathcal{B}}$, first the Multi-resolution Rand Index is defined

$$MRI(\mathcal{A},\mathcal{B}) = \begin{cases} \frac{|S_{\mathcal{A},\mathcal{B}}|+|S_{\mathcal{A}^c,\mathcal{B}^c}|+|S_{\mathcal{A},\mathcal{B}^c}|}{|S_{\mathcal{A},\mathcal{B}}|+|S_{\mathcal{A}^c,\mathcal{B}^c}|+|S_{\mathcal{A},\mathcal{B}^c}|+|S_{\mathcal{A}^c,\mathcal{B}}|} = 1 - \frac{|S_{\mathcal{A}^c,\mathcal{B}}|}{\binom{n}{2}} & \text{if } K_{\mathcal{A}} < K_{\mathcal{B}} \\ 1 - \frac{|S_{\mathcal{A},\mathcal{B}^c}|+|S_{\mathcal{A}^c,\mathcal{B}}|}{2\binom{n}{2}} & \text{if } K_{\mathcal{A}} = K_{\mathcal{B}}. \end{cases} \tag{21}$$

Note that here only set $S_{\mathcal{A}^c,\mathcal{B}}$ is responsible for the difference between clustering regardless of resolutions and set $S_{\mathcal{A},\mathcal{B}^c}$ is excluded from error quantification given the difference in resolution, where a more coarse clustering is expected when clustered at a lower resolution.

The MRI is further adjusted to produce Adjusted-MRI assuming the permutation model by

$$\begin{aligned} AMRI(\mathcal{A},\mathcal{B}) &= \frac{MRI(\mathcal{A},\mathcal{B}) - \mathbb{E}_{\text{perm}}[MRI(\mathcal{A},\mathcal{B})]}{1.0 - \mathbb{E}_{\text{perm}}[MRI(\mathcal{A},\mathcal{B})]}, \\ \mathbb{E}_{\text{perm}}[AMRI(\mathcal{A},\mathcal{B})] &= \begin{cases} 1 - \left(1 - \frac{\sum_i \binom{|A_i|}{2}}{\binom{N}{2}}\right) \frac{\sum_j \binom{|B_j|}{2}}{\binom{N}{2}} & \text{if } K_{\mathcal{A}} < K_{\mathcal{B}} \\ 1 - \frac{1}{2}\left[\left(1 - \frac{\sum_i \binom{|A_i|}{2}}{\binom{n}{2}}\right) \frac{\sum_j \binom{|B_j|}{2}}{\binom{n}{2}} + \frac{\sum_i \binom{|A_i|}{2}}{\binom{n}{2}} \left(1 - \frac{\sum_j \binom{|B_j|}{2}}{\binom{n}{2}}\right)\right] & \text{if } K_{\mathcal{A}} = K_{\mathcal{B}}. \end{cases} \end{aligned} \tag{22}$$

For comparison, we evaluate ARI and AMRI under a few simplified scenarios as described in Table S6.

### 3 Simulated data

**Producing scRNA-seq data** The SymSim package[13] was used to simulate scRNA-seq data given a known tree structure. The SymSym simulation algorithm has three knobs. The first knob simulates the within-group variation following a two-state kinetic model, in which the promoter switches between an on and an off state with certain probabilities. The second knob allows the user to simulate multiple different cell types/states, with the desired cell state structure supplied as a tree by the user. This is achieved by setting different extrinsic variability factors (EVF) for different cells assigned with the position along the tree, so that the expected absolute difference between the value of an EVF of any two cells is linearly proportional to the square root of their distance in the tree. The third knob controls the technical variation, including noise, reduced sensitivity and bias that is introduced during sample processing steps such as mRNA capture, reverse transcription, PCR amplification, RNA fragmentation, and sequencing, by simulating the major steps in experimental procedures. The separation between clusters is governed by two sets of parameters, namely the branch lengths of the tree and the within leaf standard deviation of EVF (sigma). More specifically, for any two positions in the tree, the ratio of square root tree distance to sigma determines the separability between the respective distributions of the value assigned to any given Diff-EVF.

The SymSim parameters are estimated from the mouse brain data[12]. To mitigate the effect of low sensitivity, the data is preprocessed by imputation with scVI and MAGIC. To reduce the effects of extrinsic variation, the parameters are estimated separately in each of the three largest clusters in the cortex dataset. We mainly considered two parameter settings regarding the choice of sigma, namely easier clustering tasks with sigma= 0.6 where the within-group variance is smaller, and challenging clustering tasks with sigma= 0.8 so the within-group variance is significantly higher.

**Clustering methods for simulations** *(i)* Seurat embeds an unsupervised clustering algorithm, combining dimension reduction with graph-based partitioning methods. The clusters are obtained through standard Seurat pre-processing and clustering pipeline provided by Seurat R package[8], using a range of clustering resolution from 0.05 to 3. No gene selection is performed. *(ii)* SC3 adopts consensus clustering, which summarizes clusterings from various dimension reduction procedures in a co-classification matrix as input for the final hierarchical clustering. The expressions are log normalized, and then the clustering is performed using SC3 R package[4] with $K$ varying from 2 to 12. *(iii)* UMAP followed by K-means clustering first projects data onto a low dimensional subspace by UMAP algorithm, followed by applying K-means clustering on the data in low dimensional space. Here, we use the UMAP R package with default parameters to reduce normalized expression data into three dimensions. The number of clusters is again set to be a range between 2 and 12. *(iv)* SOUP is a soft clustering method developed for scRNA-seq data, which implemented a unique pure node selection procedure. We use SOUP R Package[15] and clustering is performed for a range of $K$ from 2 to 12 with default parameters.

### 4 Single RNA-seq data analysis

#### 4.1 Data preprocessing and clustering

The preprocessing of mouse brain data started with 16,450 genes that express in at least 10 cells. We selected 2818 highly variable genes using DESCEND and SPCA (provided by '`selectGenes`' function from SOUPR R package). The counts were further depth-normalized and log-transformed to generate the input for SOUP clustering. We applied SOUP clustering for each $K$ from $K = 2$ to $K = 12$. The parameter `pure.prop` was set to 0.8 given the cells are mature. Other clustering parameters were chosen by default.

The smaller data set from human pancreas islet contains the expressions of 19950 genes of 635 cells. The counts were first depth normalized and log-transformed. Then we used the wrapper provided by SC3 R package to perform gene selection and clustering with the number of clusters ranging from 2 to 15, where one clustering is generated for each $K$. Other parameters are set all by default. The larger scRNA-seq data from five different technologies were downloaded from SeuratData R package[5]. We followed the online tutorial for the integration process and set the dimension for PCA to 40. This resulted in one integrated assay. 2000 highly variable genes were selected using '`MeanVarPlot`' method shared across data sets. They were used for the batch removal and subsequent clustering analysis. The Louvain clustering was then performed in the PCA reduced space. We sampled 50 resolution parameters within the range $[0.001, 2]$ using the Event Sampling method.

For the human brain data, the raw counts of 33,976 cells on 35,543 genes were first depth normalized and log-transformed, and each gene was mean-centered and variance-scaled. Then the effective number of UMI, donor, and library preparation batch was removed from the normalized data using a linear model. Subsequently, a set of 3715 highly variable genes were selected using 'MeanVarPlot' method and used for clustering analysis and visualization. Prior to clustering analysis, PCA was used to reduce the dimensionality of the dataset to the top 40 PCs. Then the clustering was performed using graph-based clustering implemented by Seurat with resolution parameters ranging from 0.001 to 2.

To generate an agglomerative hierarchical cluster tree from SC3, we used the `sc3` from SC3 R packages. The consensus tree was generated as a side product in the process of SC3 clustering, and we extracted it from the metadata of the SC3 object. To obtain the hierarchical tree from Seurat, we utilized the built-in function '`BuildClusterTree`' from the Seurat R package. It built an agglomerative hierarchical tree using the average distance starting from leaf nodes. The leaf nodes were the Seurat clusters generated from the highest resolution provided.

#### 4.2 More discussions for pancreas islet cells data

The uncovered hierarchical tree on 635 cells from Wang et al.[11] shares a similar structure with the tree constructed by a previous study using bimodal divisive hierarchical clustering (CellBIC[3]). The only difference is that MRtree grouped the mesenchyme cells with $\alpha$, $\beta$, $\gamma$, and $\delta$ cells in the first split, while CellBIC grouped mesenchyme cells with acinar and ductal cells. We also compared with the tree structure from SC3 agglomerative hierarchical clustering trimmed to top layers (Figure S7D). We observed that the SC3 constructed tree produced some uninterpretable hierarchical structures. For instance, two of the $\alpha$ subclusters were grouped in the same branch with acinar and ductal cells. The ductal cells were not grouped with other exocrine cells. MRtree outperformed CellBIC as well as SC3 agglomerative hierarchical clustering in that it successfully differentiated the rare delta group from beta cells.

We further applied MRtree with SC3 clusterings ($K$ from 2 to 15) on the subset of all beta cells to investigate the transcriptional heterogeneity within $\beta$ cells. We trimmed the tree at $K = 5$ to obtain five stable clusters. It shows that cluster 2 and 3 are mainly composed of cells from individuals with type 2 diabetes. Among three clusters composed of control individuals, cluster 4 are mainly from children while cluster 1 and 5 are from adults (Figure S7 E-G). In comparison, CellBIC also revealed a similar disease and age-explained division among beta cells[3]. However, their results were presented as two two-leaf trees explaining the age and disease associations, instead of a uniform hierarchical tree.

#### 4.3 More discussions for human fetal brain data

**ExN** The normalized mean expression of marker genes for upper layer neurons (i.e. *LHX2, CUX2, CUX1, SATB2, BHLHE22*) and deep layer neurons (i.e. *RORB, ETV1, FOXP1, FEZF2, TBR1, FOXP2*) and shared layer markers (i.e. *RORB, TLE4, LMO3, CRYM, ST18*) indicates gradually increased expression of markers of upper layer excitatory neurons in contrast to no expression of deep layer neuronal programs (Figure S10).

**ExM** The normalized mean expression of marker genes for upper layer neurons (i.e. *LHX2, CUX2, CUX1, SATB2, BHLHE22*) and deep layer neurons (i.e. *RORB, ETV1, FOXP1, FEZF2, TBR1, FOXP2*) and shared layer markers (i.e. *RORB, TLE4, LMO3, CRYM, ST18*) illustrates gradually increased expression of upper layer markers from ExM_0 to ExM_1 to ExM_2 in contrast to ExM_3 that expresses deep layer markers indication of layer 4/5 excitatory neurons (i.e. *LMO3, FOXP1, RORB, THY1*) (Figure S11).

**InMGE** Through the visualization of mean expression of genes that go up (i.e. *MEF2C, MAFB, SATB1, NRXN1, APP, SNAP25, YWHAB, YWHAZ, GAP43, SCN2A*) or those that go down (i.e. *MAF, DLX5, DLX2, DLX6, SYNE2*) across interneuron maturation, we observed that MGE_0 through MGE_2 follow a general maturation of MGE interneurons, in that they exhibit a gradual decrease in expression of *DLX2,-5,-6* and *MAF* transcription factors along with a gradual increase in *GAD1/2*, *LHX6*, and *SOX6* simultaneous with synaptic and axonal-related genes (i.e. *RIMS1, NRXN1, APP, GAP43*). We also observed interneuron subtype-specific genes within the most mature MGE clusters. MGE_3-MGE_5 clusters are engaged in terminal specification as such they are highly engaged in synaptic programs coincident with subtype-specification markers *SST*, *NPY*,

*NOS1, TAC1* and transcription factors like *SATB1, DLX1, SOX6, MEF2C* allowing us to divide the most mature interneurons into MGE_5 as *SST+, NPY+, NOS1+, CHODL+* interneurons and MGE_4 into a nascent *PV+* cell through expression of *TAC1+*, *MEF2C+*, and *MAFB+*. MGE_3 is slightly behind in maturation compared to MGE_4 and MGE_5 but its gene expression pattern is most closely related to MGE_4 and thus likely another *PV+* nascent interneuron type (perhaps closely related to the *PV+* cell type found in Tasic, 2016[9] which co-express *MEIS2*) (Figure S12A). The significant PPl networks exhibit different and more complex patterns as the cluster is more mature. For example, the MGE_2 network, while not significant (Table S7), illustrates simple direct interactions between structural cytoskeleton genes, showing that these cells are heavily engaged in growth, whereas MGE_1 network, again while not significant, depicts higher engagement of canonical interneuron *DLX* transcription factors, which reduce expression in later developmental stages such as in MGE_0 whose significant network involves transcription factors *MAF* and *MAFB*, as well as AMPA-receptor and pre-synaptic function illustrating a gradual increase in function and shifting canonical interneuron transcriptional programs across maturation (Figure S12B-D). The other MGE cluster significant PPI's depict nascent subtypes of MGE-derived interneurons. InMGE_5 PPI network is more complex, including many directly connected genes related to axonal and synaptic development (i.e., post-synaptic GABAergic machinery, *APP, GAP43, NMDA*-receptor components) as well as subtype-specific genes such as NOS1. MGE_3 PPI depicts a more complex network elaborating on APP, GABAergic post-synaptic components found in MGE_5 and illustrating unique direct interactions of *CALM1, RBFOX1, SEMA and EPHN, CAMK4* that show these cells are not only more mature but also very different from MGE_0, MGE_1, MGE_2 and MGE_5. Interestingly, MGE_3 PPI is more similar to MGE_4 but expresses similar genes to a canonical excitatory neuron lineage in select aspects such as *LHX2, PAX6, MEIS2* that maybe be very similar to specific MGE cell-type found in Tasic, 2016[9]. MGE_4 PPI network is very similar to MGE_3 with more genes added to *CALM1* and *APP* connected network genes including presynaptic components, *NMDA*-receptor components and action potential related machinery owing to their more mature functional state (Figure S12E-G).

| Method | method | optimal K |
|---|---|---|
| SC3 | $K_{TW} - 1$ based on TW distribution | 22 |
| SOUP | Cross validation | 13(11) |
| tSNE + K-Means | Adaptive density peak detection (ADPclust) | 6 |
| PCA + K-Means | K that achieved maximum Gap statistics | 9 |

Table S1: The optimal number of clusters determined by different $K$-selection methods.

**InCGE** CGE_0 through CGE_2 follow a general maturation of CGE interneurons through gradual decrease in expression of *DLX2,-5,-6* and *ARX* transcription factors along with a gradual increase in *GAD1/2, NR2F1, NR2F2, PROX1* simultaneous with synaptic and axonal-related genes (i.e. *RIMS1, NRXN1, GAP43, MEF2C*). While all CGE clusters show expression for subtype marker *CALB2*, only the most mature cluster, CGE_2, expresses the subtype marker *VIP+* and *CCK+*. Given this time in development, these states fit nicely with the fact that CGE interneurons are born after MGE interneurons, and while both cell types are born in the ventral telencephalon, their terminal specification happens only upon beginning synaptogenesis when they begin to express subtype-specific markers (Figure S13B). Significant protein-protein interacting (PPI) of CGE_0 illustrates the presence of *DLX2* and *DLX5* but also that this cluster is heavily engaged in translation with similar ribosomal gene network found within intermediate progenitor (IP) cells IP_0 and to the least mature deep excitatory neurons ExDP_2 that also express ribosomal and IP-specific genes. Interestingly this PPI also highlights *BTG1*, a factor known to anti-regulate cell-cycling and promote neuronal lineages. Taken together, this cluster is likely a CGE-derived IP-like interneuron. PPI network from CGE_1 shows only two connected genes *KIF21A* and *TPR* involved in mRNA transport thus this represents a small step forward in maturation. While PPI networks formed from CGE_2 also only have two connected genes *GRIK2* and *GRIN2B* critically involved in post-synaptic glutamate signaling and plasticity, thus in combination with gene expression patterns showing these cells are the most mature *VIP+ CCK+ CR+* CGE-derived interneurons (Figure S13C-E).

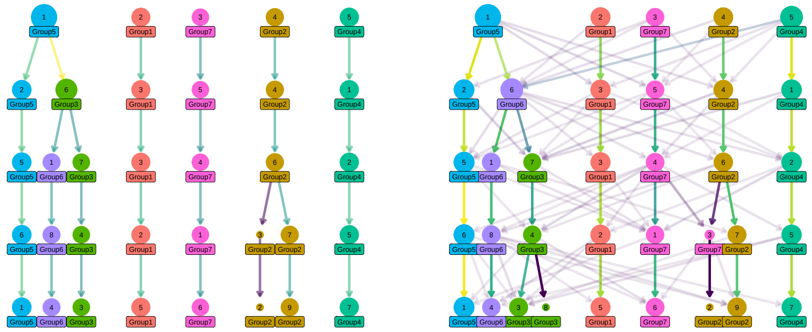

Figure S1: Right: true hierarchical cluster tree underlying the simulated single-cell data from five clusters at the top layer to eight clusters at the bottom layer. Right: the observed noisy cluster tree obtained from applying flat clustering algorithm with increasing resolutions on the simulated data. The goal of MRtree is to recover the underlying hierarchical cluster tree from the noisy cluster tree.

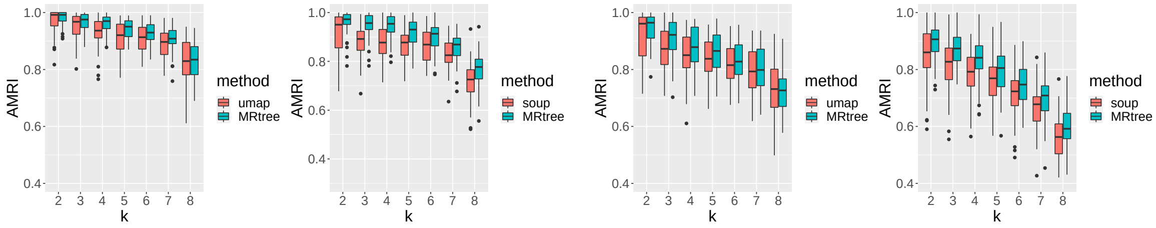

(a) easy scenario

(b) challenging scenario

Figure S2: Evaluation of MRtree on clustering accuracy with simulated data. Comparing the accuracy of MRtree clusters with the initial flat clustering results obtained at multiple resolutions using SOUP and UMAP+Kmeans, under an easy scenario with smaller within-group variance (a), and a challenging scenario with higher within-cluster variance (b). The accuracy is measured by AMRI.

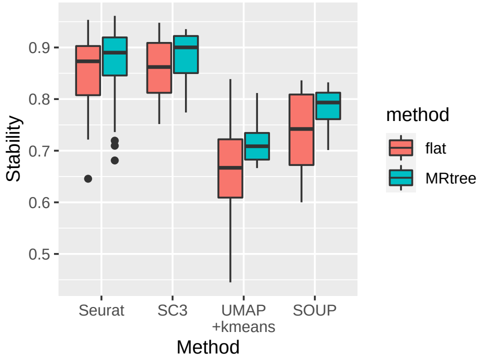

Figure S3: Cluster stability for MRtree clusters versus flat clustering results on simulated data. Comparisons are conducted at the resolution corresponding to the actual number of clusters to avoid confounding effects. The results for repeated experiments are shown by the boxplot.

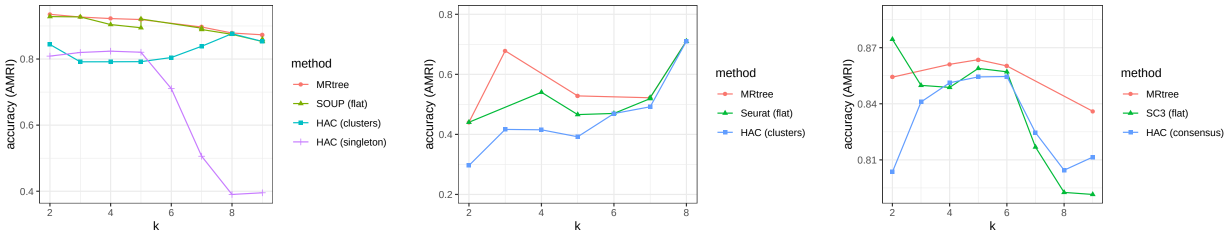

(a) SOUP (b) Seurat (c) SC3

Figure S4: Clustering accuracy for mouse brain data. Comparing accuracy measure at each resolution for MRtree constructed tree, initial flat clustering, and agglomerative hierarchical clustering using SOUP (a), Seurat (b), and SC3 (c).

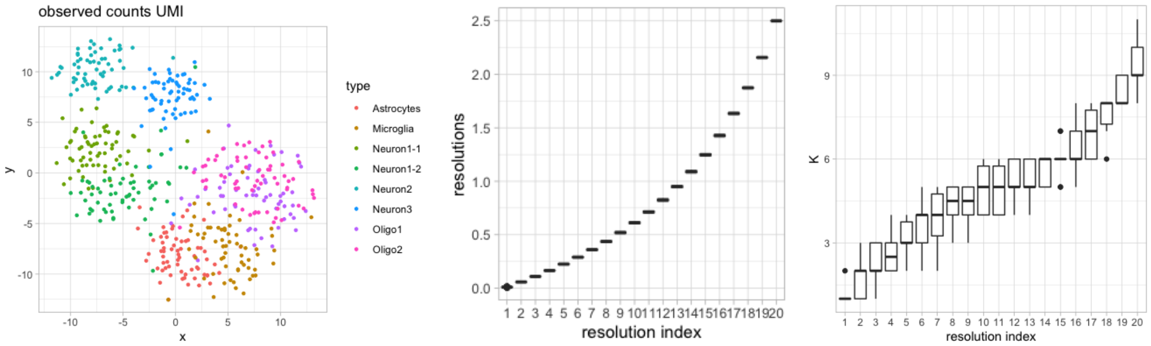

Figure S5: Evaluation of Event Sampling on simulated data. Left: UMAP projection of the simulated data belonging to 8 clusters. Middle: the sampled resolution parameters versus sample index. Right: the number of resulting clusters using Seurat clustering from the sampled resolution parameters for each resolution index, respectively.

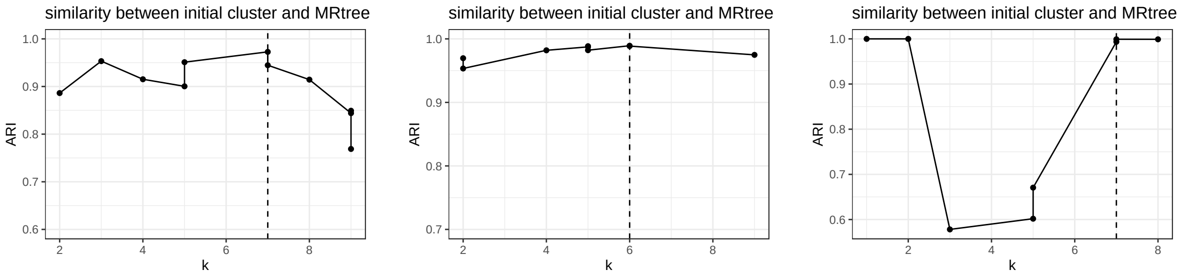

(a) SOUP (b) SC3 (c) Seurat

Figure S6: Clustering stability analysis of mouse brain data, derived by comparing the reconciled clustering results with the initial flat clusterings measured by ARI from SOUP (a), SC3 (b) and Seurat (c).

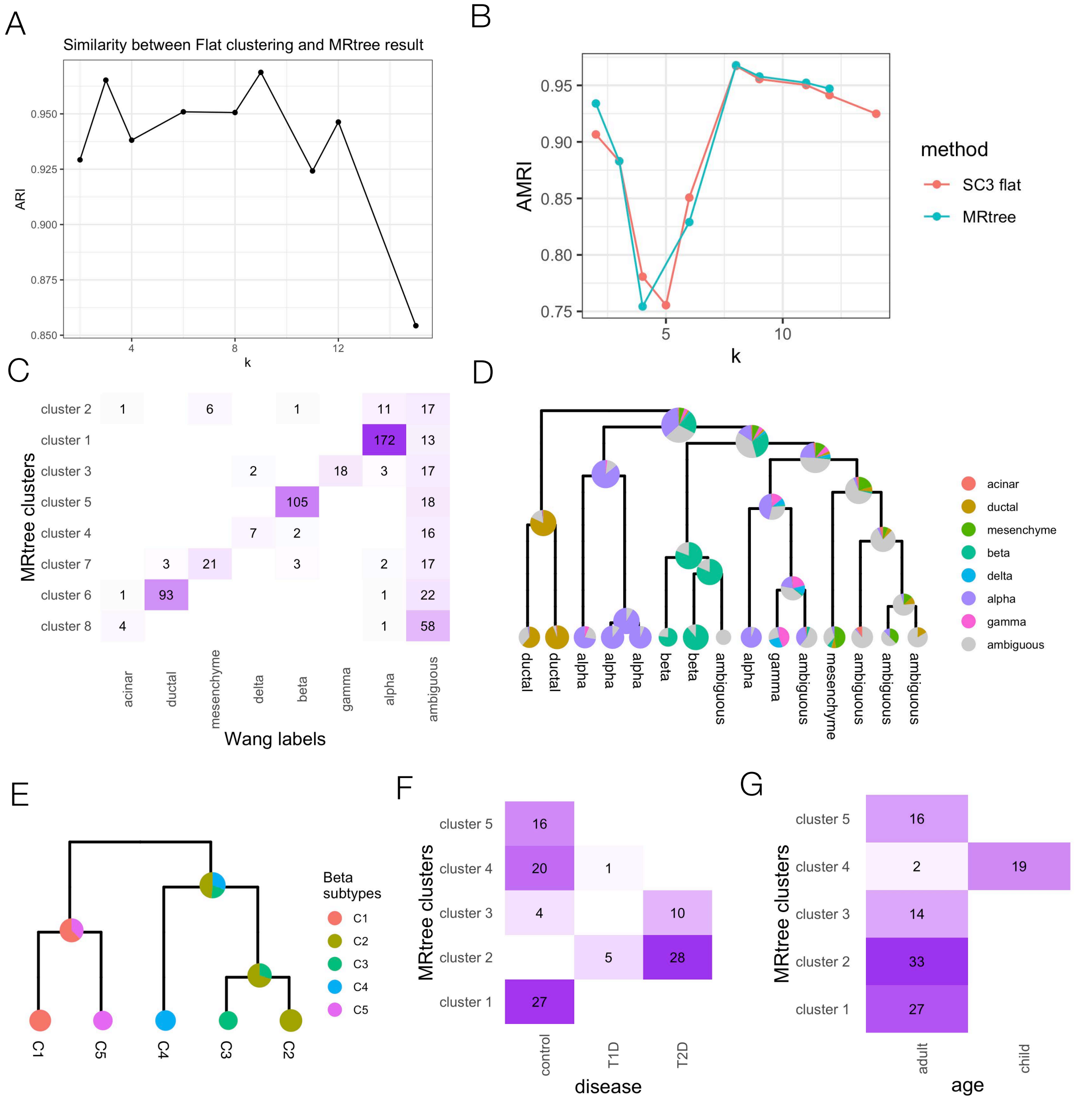

Figure S7: MRtree applied on 635 pancreas islet cells from Wang et al.[11]. (A) Stability plot from the MRtree results. ARI between the initial SC3 flat clusterings and layers of clusterings in MRtree constructed tree. Higher ARI indicates the initial flat clustering is close to the reconciled MRtree clusters and thus stable. (B) Clustering accuracy calculated for each resolution (number of resulting clusters, $K$) comparing the initial SC3 clusters (red) and MRtree clusters (blue). (C) Heatmap of the confusion matrix comparing the MRtree clusters obtained at $K = 8$ with the Wang labels. (D) Hierarchical cluster tree constructed via SC3 agglomerative hierarchical clustering, starting with one cell per cluster, trimmed to top 15 layers. Leaf labels show to which cluster the majority of cells belong. (E) The MRtree-constructed tree for 111 $\beta$ cells, trimmed to five leaf-clusters with color showing MRtree subtype labels. (F-G) Examine the five MRtree $\beta$ subtypes with respect to disease (F) and age (G) by visualizing the count matrix. The value in each entry indicates the number of cells in the corresponding cluster associated with the attribute.

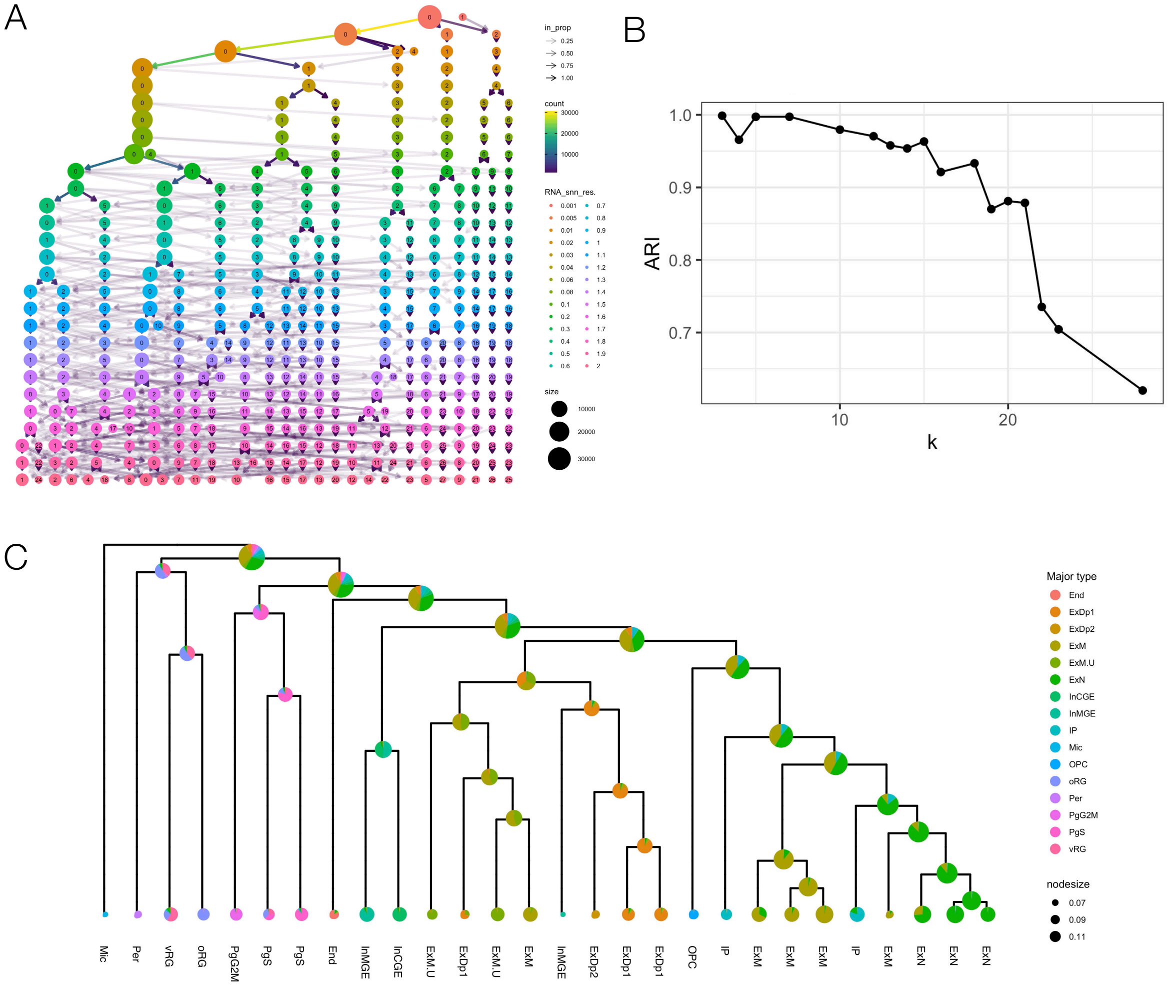

Figure S8: MRtree and Seurat hierarchical clustering results on cells from human brain data. (A) Original cluster tree formed by flat clusterings via Seurat with a range of resolutions from 0.001 to 2. (B) The similarity measured by Adjusted Multiresolution Rand Index (AMRI), between each layer of initial clusters from flat clustering and constructed hierarchical clusterings as a measure of clustering stability. (C) Dendrogram showing the hierarchy generated by agglomerative hierarchical clustering starting with Seurat clusters obtained at a high resolution (resolution parameters set to 2).

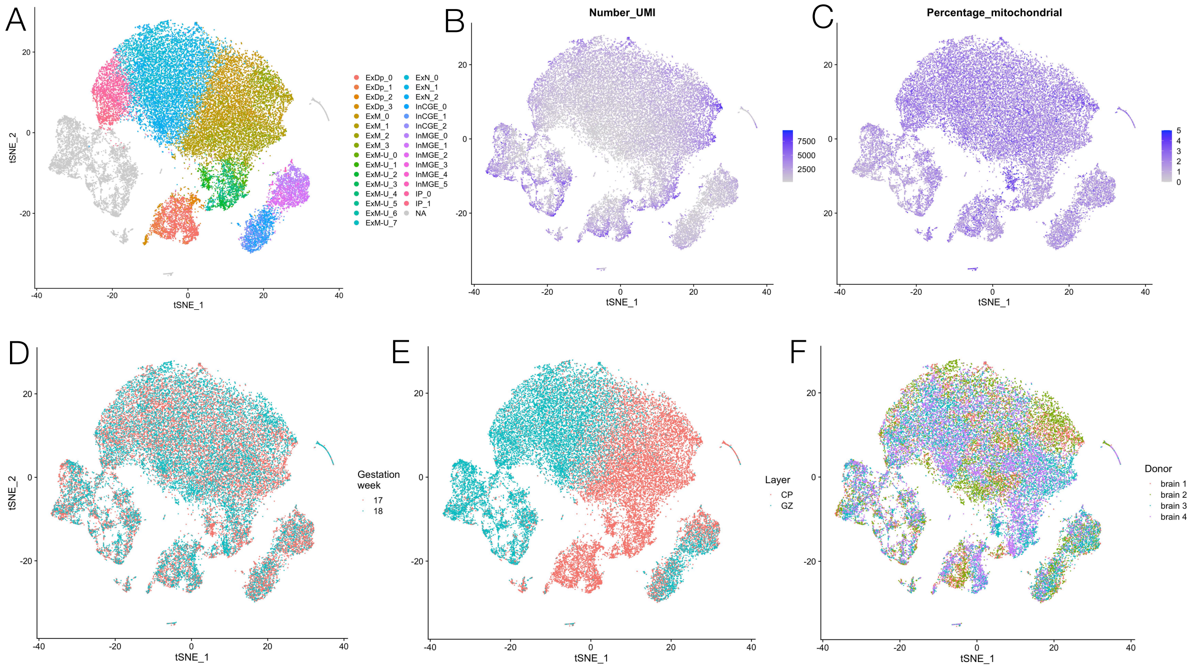

Figure S9: MRtree sub clustering results were driven by biological, not technical variation. (A) tSNE projection of MRtree sub-clustering of neurons in human brain data. (B) tSNE projection of the mean UMI (unique molecular identifier) counts per cell. (C) tSNE projection of the mean percentage of mitochondria reads per cell counts. (D) tSNE projection of the gestational week (GW) of each cell tissue origin. (E) tSNE projection of the cortical region, either cortical plate (CP) or germinal zone (GZ) from which cells were derived from. (F) tSNE projection of the brain sample from which each cell is derived from.

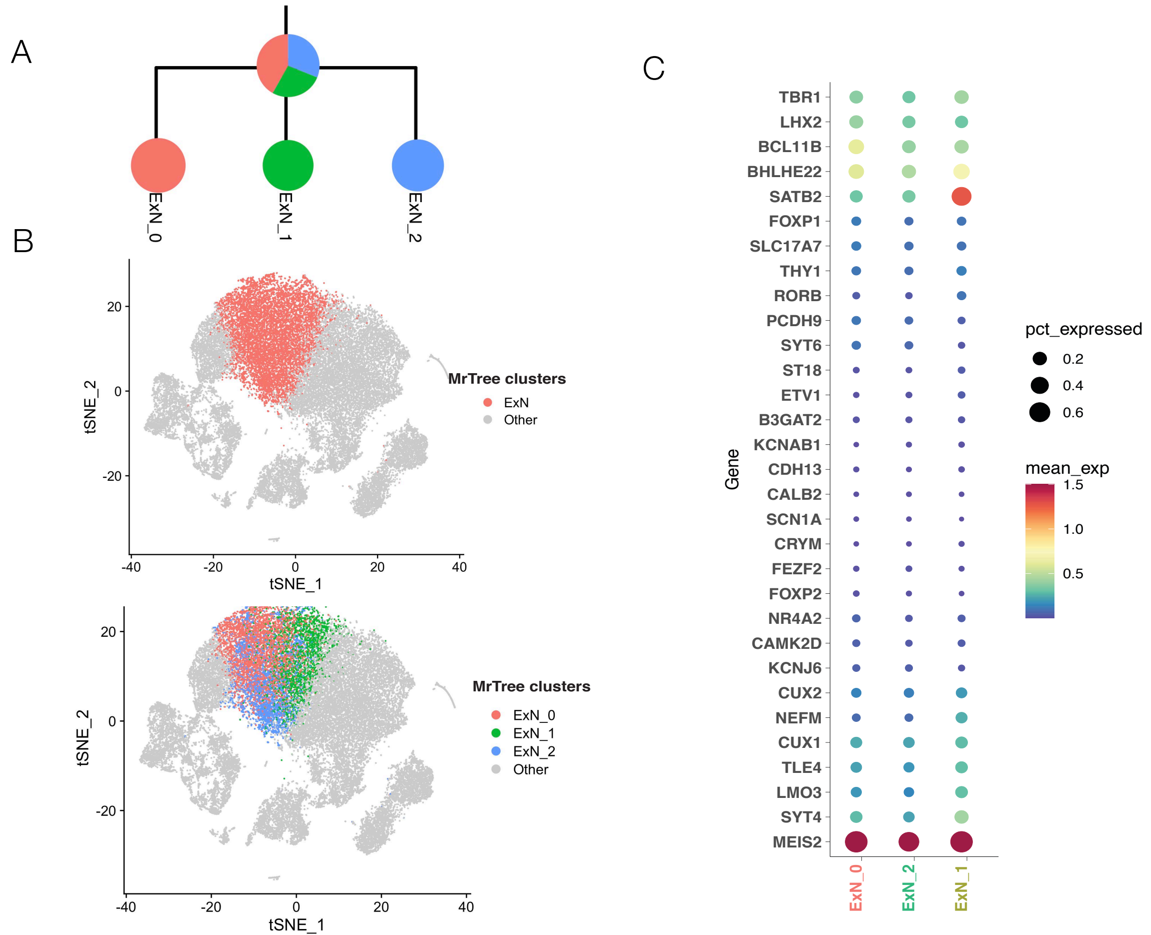

Figure S10: ExN migrating neurons follow the developmental trajectory of migrating upper layer excitatory neurons. (A) Hierarchical cluster tree showing the subtype structure of migrating excitatory neurons (ExN) generated by MRtree from initial Seurat multiresolution clustering, resulting in three stable subclusters. (B) top: tSNE plot of all cells where ExN identified by MRtree are colored by red, bottom: tSNE plot where ExN are broken into ExN_0, colored in red, ExN_1, colored by green, and ExN_2 colored in blue. (C) Gene expression dot plot showing the normalized mean expression of marker genes for upper-layer neurons (i.e. *LHX2, CUX2, CUX1, SATB2, BHLHE22*) and deep layer neurons (i.e. *RORB, ETV1, FOXP1, FEZF2, TBR1, FOXP2*) and shared layer markers (i.e. *RORB, TLE4, LMO3, CRYM, ST18*).

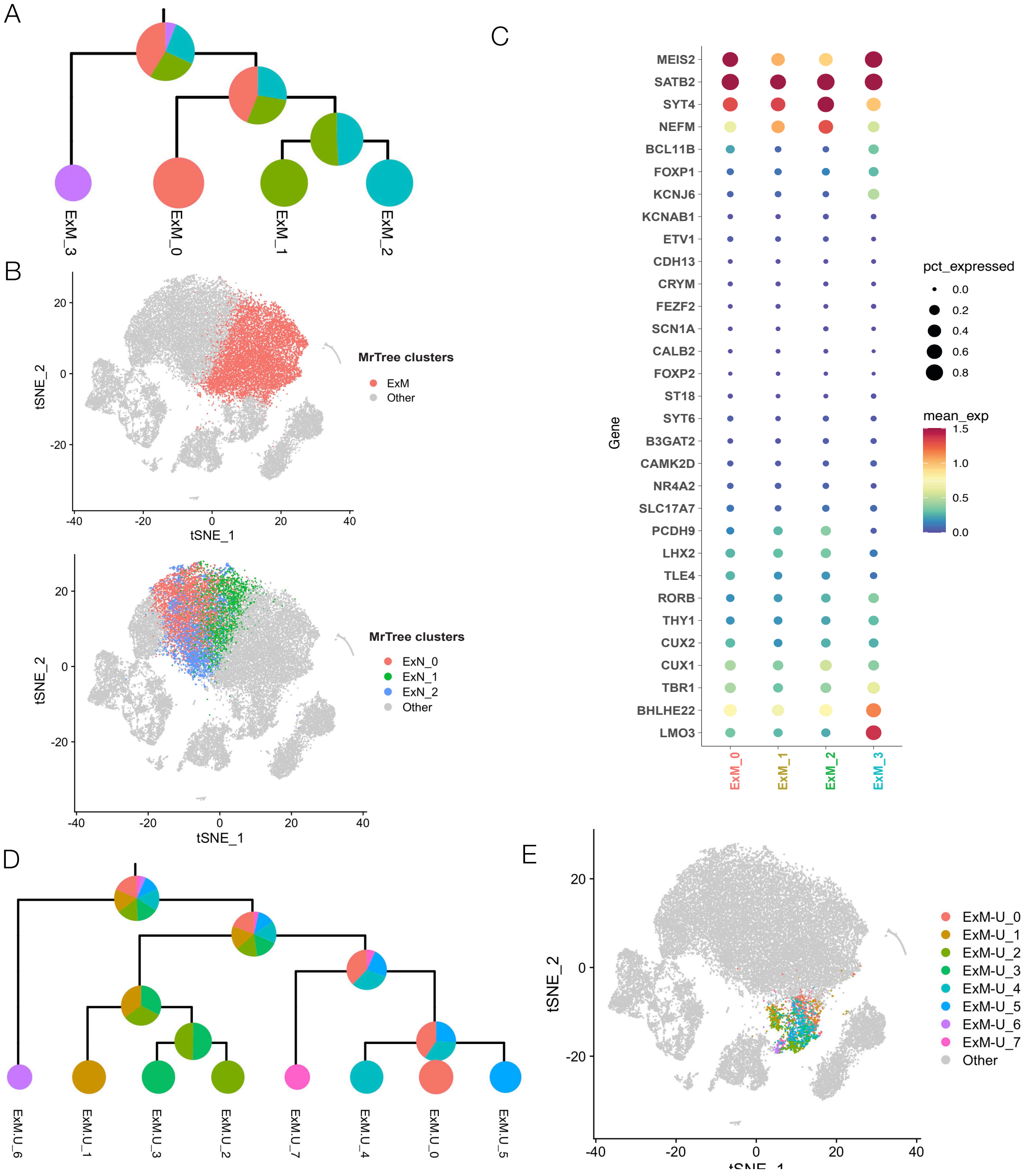

Figure S11: MRtree subtypes of ExM maturing excitatory neuron and upper layer enriched neurons. (A) Hierarchical cluster tree showing the subtype structure of maturing excitatory neurons (ExM) generated by MRtree from initial Seurat multiresolution clustering, resulting in four stable subclusters. (B) top: tSNE projection with ExM colored by red, bottom: tSNE projection of MRtree clustering where ExM is broken into ExM_0, colored in red, ExM_1 colored by green, and ExM_2 colored in blue, and ExM_3 colored in purple. (C) Gene expression dot plot showing the normalized mean expression of marker genes for upper-layer neurons (i.e. *LHX2, CUX2, CUX1, SATB2, BHLHE22*) and deep layer neurons (i.e. *RORB, ETV1, FOXP1, FEZF2, TBR1, FOXP2*) and shared layer markers (i.e. *RORB, TLE4, LMO3, CRYM, ST18*). (D) Hierarchical cluster tree showing the subtype structure of upper layer migrating excitatory neurons (ExM-U) generated by MRtree, which results in eight subclusters.(E) tSNE projection with ExM-U cells colored according to the identified eight subtypes.

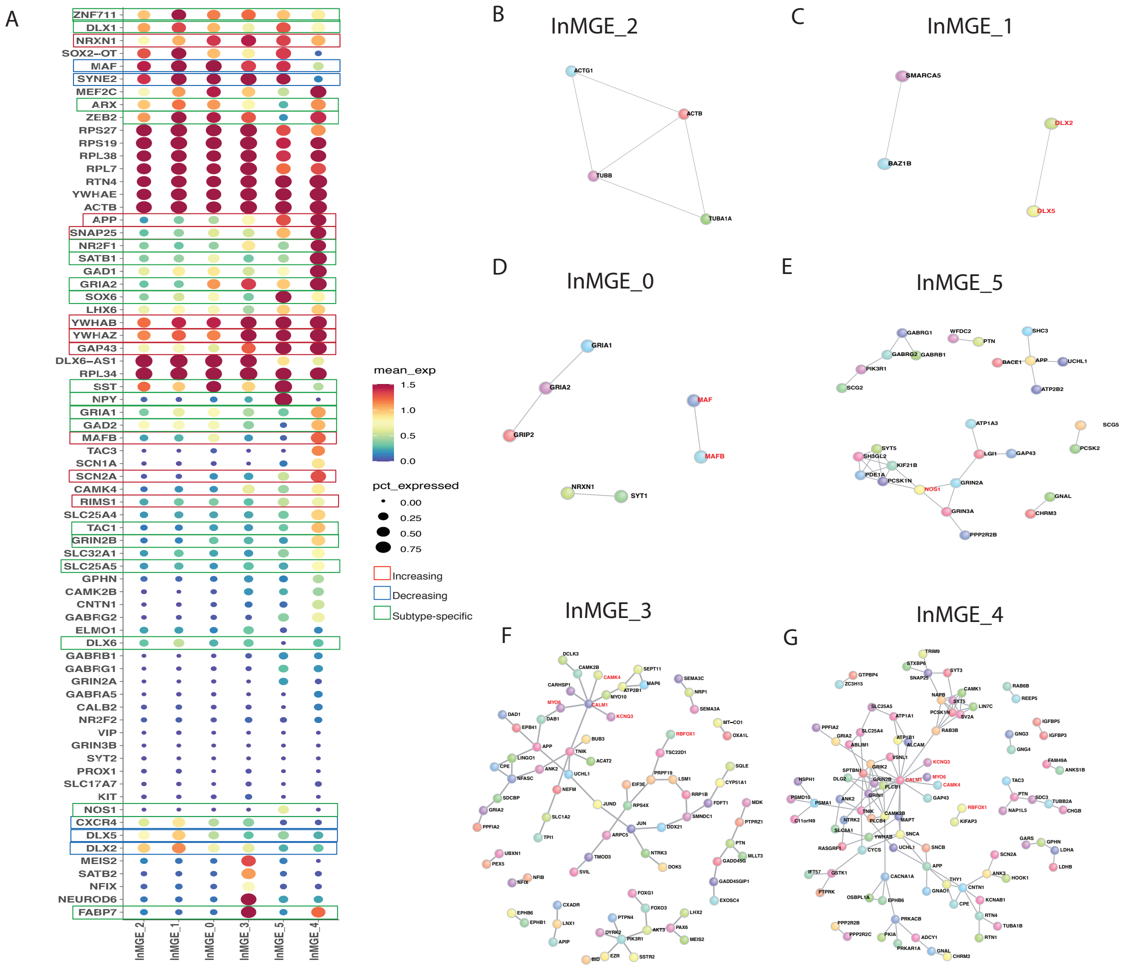

Figure S12: MGE Interneurons are split by cell type and maturation state by MRtree. (A) Gene expression dot plot showing the normalized mean expression of transcription factors and genes indicative of interneuron development that go up across maturation in red brackets (i.e., *MEF2C, MAFB, SATB1, NRXN1, APP, SNAP25, YWHAB, YWHAZ, GAP43, SCN2A*) and those that go down across interneuron development in blue brackets (i.e., *MAF, DLX5, DLX2, DLX6, SYNE2*) and select sub-type specific genes in green brackets (i.e., *SATB1, SST, NOS1, NPY, TAC3, TAC1, SLC25A5, MEF2C*) across all MGE-derived interneuron clusters. (B-G) Protein-protein interacting (PPI) networks formed from differential genes (DGE) expressed in B) MGE_2*, C) MGE_1*, D) MGE_0, E) MGE_5, F) MGE_3 and G) MGE_4, arranged by maturation.

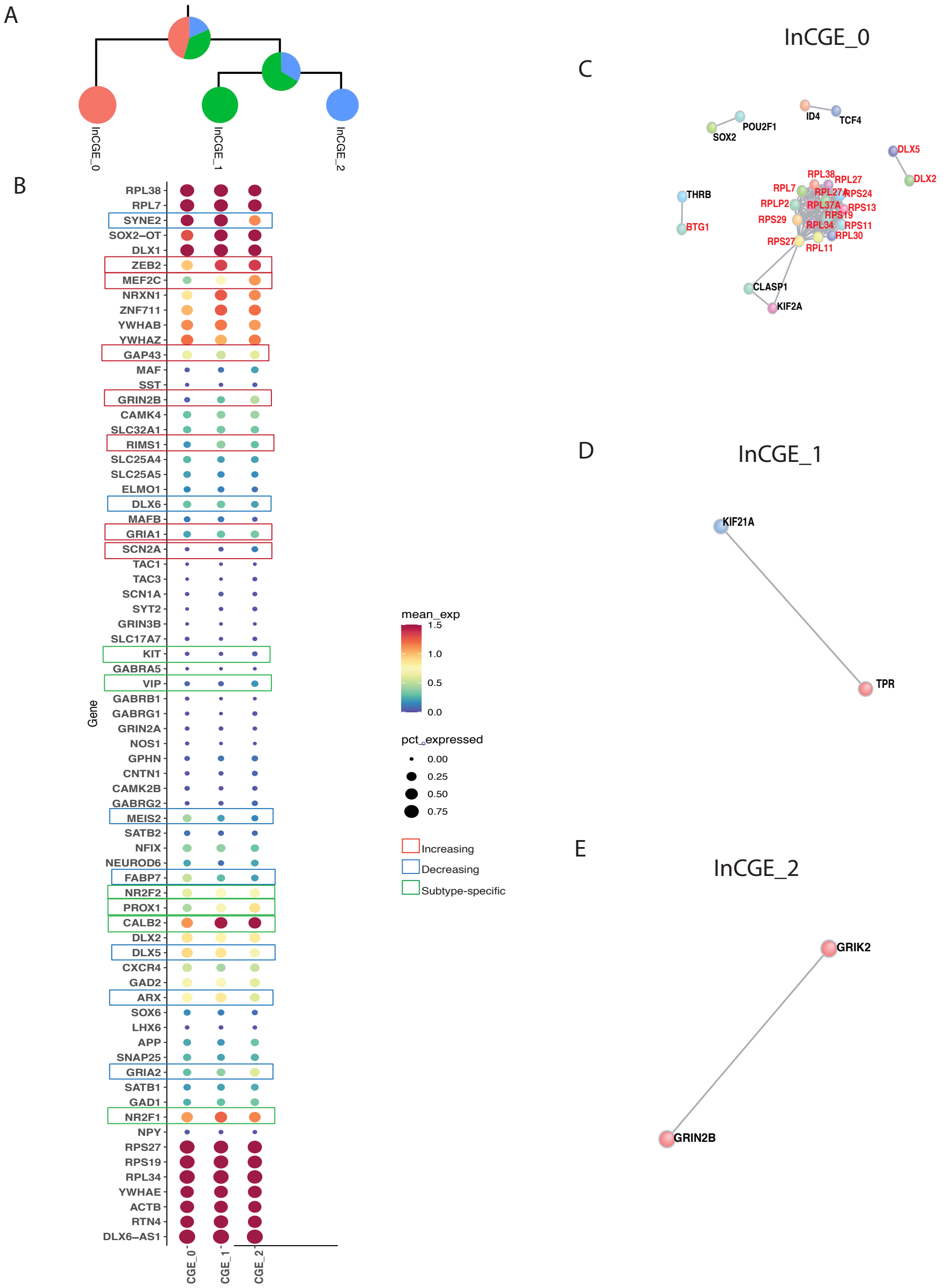

Figure S13: CGE Interneurons are split by subtle changes in maturational state at high resolution. (A) Hierarchical cluster tree showing the subtype structure of CGE interneurons generated by MRtree from initial Seurat multiresolution clustering, resulting in three stable subclusters. (B) Gene expression dot plot showing the normalized mean expression of transcription factors and genes indicative of interneuron development that go up across maturation in red brackets (i.e., *ZEB2, NR2F1, NR2F2, GRIN2B, NRXN1, RIMS1 GRIA1, GAP43*) and those that go down across interneuron development in blue brackets (i.e., *DLX5, DLX2, DLX6, ARX, SYNE2*) and select sub-type specific genes in green brackets (i.e., *VIP, CALB2 (Calretinin), PROX1*) across all CGE-derived interneuron clusters. (C-E) Significant protein-protein interacting (PPI) networks formed from differential genes (DGE) expressed in CGE_0 (C), CGE_1 (D), and CGE_2(E).

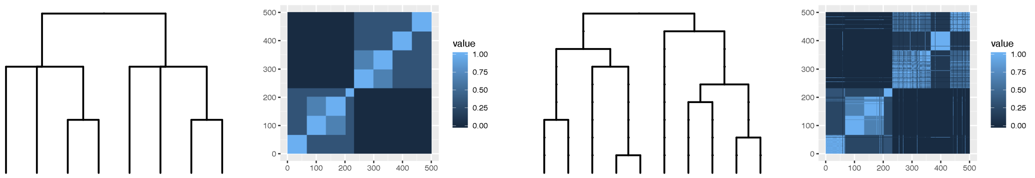

(a) True tree (b) Recovered tree

Figure S14: Hierarchical cluster tree visualized by dendrograms and their corresponding similarity matrices. The entry of the similarity matrix is the length of the shared branch in the dendrogram. (a) The true hierarchy that represents cluster similarity; (b) the recovered tree structure via tree-construction methods.

| Splits | Cell group | Marker genes |
|---|---|---|
| (acinar, ductal) from (mesenchyme, $\delta$ , $\beta$, $\alpha$, $\gamma$) | (acinar, ductal) | *SPP1, REG1A, REG1B, PRSS1, CRP, SERPINA3, MMP7, PIGR, REG3A, LCN2, TRY6, CFB, OLFM4, CFTR, C3* |
| | (mesenchyme, $\delta$ , $\beta$, $\alpha$, $\gamma$) | *IAPP, INS-IGF2, SCG2, CHGB, GCG, INS, SCG5, CPE, PPY, SLC30A8, COL1A2, PAX6, ERO1LB, TTR, ENPP2* |
| mesenchyme from ($\delta$ , $\beta$, $\alpha$, $\gamma$) | mesenchyme | *FN1, COL1A2, COL6A3, COL1A1, TIMP3, COL3A1, LUM, IGFBP5, AXL, TIMP1, ACTA2, SPARC, BMP8A, TNFAIP6, VCAN* |
| | ($\delta$ , $\beta$, $\alpha$, $\gamma$) | *TTR, CHGB, SLC30A8, TM4SF4, ALDH1A1, SCG5, CPE, GCG, G6PC2, PDK4, SCGN, PAX6, CLU, SCG3, GC* |
| new cluster from ($\delta$ , $\beta$, $\alpha$, $\gamma$) | new cluster | *SPARC, STC1, PECAM1, COL4A1, A2M, RNASE1, CD34, NFIB, PRDM1, ESM1, THBS1, CD93, FLT1, EMCN, CXCR4* |
| | ($\delta$ , $\beta$, $\alpha$, $\gamma$) | *G6PC2, SCGN, BBIP1, PTEN, DSP, FAM46C, PTPRN, SYT4, PCSK2, SNAP25, TLK1, CADM1, CDCP1, ENPP2, BEX1* |
| ($\delta$ , $\beta$) from ($\alpha$, $\gamma$) | ($\delta$ , $\beta$) | *IAPP, INS-IGF2, INS, HADH, PFKFB2, ADCYAP1, RBP4, PCSK1, DLK1, SORL1, G6PC2, CASR, BMP5, NPTX2, TGFBR3* |
| | ($\alpha$, $\gamma$) | *GCG, TM4SF4, TTR, GC, PDK4, FAP, RGS4, ALDH1A1, LOXL4, MUC13, LDHA, TMEM176B, FXYD5, VIM, EZR* |

Table S2: Top 15 significant marker genes (adjusted p-value $< 0.05$) for tree splits from MRtree constructed tree on 635 human pancreas islet cells.

| Cluster | Cell type | Marker genes |
|---|---|---|
| 1 | $\alpha$ | *GCG, RGS4, TTR, GC, PDK4, TM4SF4, ALDH1A1, LOXL4, PCSK2, PLCE1, SCG2, CHGB, FAP, GPX3, SLC38A4* |
| 2 | new cluster | *PECAM1, CD34, SPARC, ESM1, CD93, FLT1, EMCN, PRDM1, KDR, CXCR4, ELTD1, MYCT1, GPR116, PODXL, GNG11* |
| 3 | $\gamma$ | *PPY, FLJ43390, THSD7A, SLITRK6, UNC80, MIR7-3HG, CALB1, SEC11C, ETV1, TMEM47, KIAA1377, CADPS, ABCC9, RMST, PRG4* |
| 4 | $\delta$ | *RBP4, CASR, HADH, PCSK1, PDX1* |
| 5 | $\beta$ | *IAPP, INS-IGF2, INS, PFKFB2, HADH, G6PC2, ADCYAP1, PCSK1, DLK1, SORL1, SCGN, BMP5, ABCC8, SLC30A8, CASR* |
| 6 | ductal | *SPP1, CRP, CEACAM7, PIGR, CFTR, MMP7, TSPAN8, TNFAIP2, APCS, CD44, ANXA4, LCN2, VCAM1, SLC4A4, CEACAM6* |
| 7 | mesenchyme | *COL6A3, COL1A2, TIMP3, COL3A1, COL1A1, FN1, IGFBP5, LUM, AXL, SPARC, TIMP1, ACTA2, BMP8A, VCAN, TNFAIP6* |
| 8 | acinar | *REG1B, PRSS1, REG1A, REG3A, TRY6, PRSS2, CPB1, CELA3A, CPA2, CTRB2, PRSS3, PNLIP, SPINK1, CTRB1, LYZ* |

Table S3: Top 15 significant marker genes (adjusted p-value $< 0.05$) for each of the eight clusters identified from MRtree on 635 human pancreas islet cells.

| Cell type | Short name | Number of subtypes | Number of cells |
|---|---|---|---|
| Radial glia | oRG | 5 | 1293 |
| | vRG | 4 | 984 |
| Cycling progenitors | PgS | 5 | 1232 |
| | PgG2M | 5 | 695 |
| Intermediate progenitors | IP | 4 | 2150 |
| Migrating excitatory neurons | ExN | 8 | 9995 |
| Maturing excitatory neurons | ExM | 8 | 9822 |
| | ExM-U | 6 | 1756 |
| Excitatory deep layer 1/2 | ExDp1 | 4 | 2039 |
| | ExDp2 | 2 | 166 |
| Interneurons | InCGE | 6 | 1434 |
| | InMGE | 8 | 1705 |
| Oligodendrocyte precursors | OPC | 6 | 306 |
| Endothelial | End | 4 | 237 |
| Pericytes | Per | 3 | 114 |
| Microglia | Mic | 1 | 48 |

Table S4: 16 major cell types identified in Polioudakis et al.[6]

| | |
|---|---|
| InMGE | SST, MAF, ERBB4, DLX6-AS1, PDZRN4, ARX, SOX2-OT, GAD1, LHX6, DLX2 |
| InCGE | DLX6-AS1, DLX1, CALB2, PDZRN3, SOX2-OT, ERBB4, PLS3, SCGN, DLX5, PROX1 |
| Per | COL4A1, RGS5, IGFBP7, SPARC, DCN, ITIH5, CALD1, NDUFA4L2, SPARCL1, IFITM3 |
| End | ITM2A, IGFBP7, CLDN5, GNG11, ARHGAP29, SLC38A5, HBG2, FN1, HLA-E, FLT1 |
| Mic | CCL3, SPP1, CCL3L3, CCL4, CCL4L2, IL1B, RGS1, AIF1, A2M, P2RY12 |
| ExDp1 | LMO3, CRYM, FBXW7, TLE4, LMO7, PPP1R1B, MEG3, KCTD12, PDE1A, GRIN2B |
| ExDp2 | LMO3, SERPINI1, GRIK2, SYT1, MEG3, SEMA3E, NRCAM, NCAM2, B3GALT2, UTRN |
| OPC | BCAS1, PDGFRA, PMP2, OLIG1, SCRG1, BCAN, APOD, S100B, LIMA1, EPN2 |
| RG | PTN, VIM, HES1, ID4, FOS, PEA15, SFRP1, EGR1, SLC1A3, CLU |
| Pg | TOP2A, HMGB2, MKI67, CENPF, NUSAP1, HIST1H4C, HIST1H1D, ASPM, SMC4, PRC1 |
| ExN | ENC1, EPHB6, ROBO2, RASD1, NRP1, HS3ST1, KCNQ3, PRKX, SDCBP, MEIS2 |
| IP | PPP1R17, SSTR2, EOMES, NHLH1, TMEM158, PENK, HES6, CORO1C, CA12, NRN1 |
| ExM | SATB2, SYT4, LIMCH1, PLXNA4, DAB1, MCUR1, MASP1, GAP43, STMN2, ARPP21 |
| ExM-U | NEFL, MEF2C, NEFM, RUNX1T1, LMO4, NPY, R3HDM1, LPL, RSPO3, LRRN3 |

Table S5: Top 10 marker genes for each of 14 major cell types, ranked by the average log fold change among the significant genes identified by the FDR-adjusted p-values.

| Modifications of $\mathcal{A}'$ | $RI(\mathcal{A}, \mathcal{A}')$ | $MRI(\mathcal{A}, \mathcal{A}')$ |
|---|---|---|
| Two clusters joined | $1 - \frac{2n}{(n-1)K^2}$ | 1 |
| One cluster split into two equal parts (even) | $1 - \frac{n}{2(n-1)K^2}$ | 1 |
| One point taken from each cluster to form a new cluster of $K$ points | $1 - \frac{K^2-K+2n-2K}{n(n-1)}$ | $1 - \frac{K^2-K+K^2-K}{n(n-1)}$ |
| $\frac{n}{K}$ clusters are formed with $K$ points in each, one point from original cluster ($K < \frac{n}{K}$) | $1 - \frac{K^2+n-2K}{K(n-1)}$ | $1 - \frac{2(K-1)}{n-1}$ |

Table S6: Compare the performance of RI and MRI under a few simplified scenarios. $\mathcal{A} = \{A_1, \ldots, A_K\}$ is a clustering of $n$ data points where each cluster is of size $\frac{n}{K}$. $\mathcal{A}'$ is a partition of the same data modified from $\mathcal{A}$ as specified in the first column.

| | IP | | ExDp | | | | MGE | | | | | | CGE | | |
|---|---|---|---|---|---|---|---|---|---|---|---|---|---|---|---|
| | 1 | 2 | 2 | 0 | 1 | 3 | 2 | 0 | 1 | 5 | 3 | 4 | 0 | 1 | 2 |
| Direct Edges Count P-Value | 0.003 | 0.000 | 0.000 | 1 | 0.000 | 0.000 | 0.138 | 0.001 | 0.128 | 0.000 | 0.000 | 0.000 | 0.000 | 0.229 | 0.004 |
| Seed Direct Degrees Mean | 0.003 | 0.007 | 0.010 | 1 | 0.002 | 0.000 | 0.043 | 0.016 | 0.495 | 0.000 | 0.127 | 0.000 | 0.000 | 0.465 | 0.081 |
| Seed Indirect Degrees Mean | 0.000 | 0.000 | 0.000 | 0.356 | 0.000 | 0.000 | 0.001 | 0.022 | 0.336 | 0.000 | 0.246 | 0.000 | 0.000 | 0.093 | 0.001 |
| CI Degrees Mean | 0.021 | 0.082 | 0.005 | 0.287 | 0.001 | 0.009 | 0.397 | 0.050 | 0.574 | 0.000 | 0.894 | 0.000 | 0.000 | 0.084 | 0.001 |
| #Direct Network Genes | 55 | 21 | 130 | 0 | 24 | 90 | 4 | 7 | 4 | 28 | 81 | 96 | 18 | 2 | 2 |
| #Direct Interactions Genes | 124 | 21 | 2309 | 0 | 22 | 136 | 5 | 4 | 2 | 31 | 76 | 131 | 107 | 1 | 1 |
| Mean Associated Protein Direct Connectivity | 4.509 | 2 | 35.52 | 0 | 1.833 | 3.022 | 2.5 | 1.142 | 1 | 2.214 | 1.876 | 2.729 | 11.88 | 1 | 1 |
| Mean Associated Protein Indirect Connectivity | 241.8 | 34.85 | 2742 | 6.941 | 30.26 | 87.46 | 394.4 | 11.33 | 6.704 | 41.66 | 52.88 | 82.16 | 1360 | 26 | 16.66 |
| Mean CI connectivity | 4.056 | 2.580 | 9.550 | 2.102 | 2.652 | 3.363 | 2.415 | 2.203 | 2.806 | 3.157 | 2.774 | 3.367 | 5.984 | 2 | 2.202 |

Table S7: Summary statistics for PPI Networks for MRtree fetal brain subtypes.

### References

1. Lawrence Hubert and Phipps Arabie. Comparing partitions. *Journal of classification*, 2(1): 193–218, 1985.

2. Lucas GS Jeub, Olaf Sporns, and Santo Fortunato. Multiresolution consensus clustering in networks. *Scientific reports*, 8(1):1–16, 2018.

3. Junil Kim, Diana E Stanescu, and Kyoung Jae Won. Cellbic: bimodality-based top-down clustering of single-cell rna sequencing data reveals hierarchical structure of the cell type. *Nucleic acids research*, 46(21):e124–e124, 2018.

4. V. Y. Kiselev, K. Kirschner, M. T. Schaub, T. Andrews, A. Yiu, T. Chandra, K. N. Natarajan, W. Reik, M. Barahona, A. R. Green, and M. Hemberg. Sc3: consensus clustering of single-cell rna-seq data. *Nature Methods*, 14(5):483–+, 2017. ISSN 1548-7091.

5. Satija Lab. *panc8.SeuratData: Eight Pancreas Datasets Across Five Technologies*, 2019. R package version 3.0.2.

6. Damon Polioudakis, Luis de la Torre-Ubieta, Justin Langerman, Andrew G Elkins, Xu Shi, Jason L Stein, Celine K Vuong, Susanne Nichterwitz, Melinda Gevorgian, Carli K Opland, et al. A single-cell transcriptomic atlas of human neocortical development during mid-gestation. *Neuron*, 103(5):785–801, 2019.

7. William M Rand. Objective criteria for the evaluation of clustering methods. *Journal of the American Statistical association*, 66(336):846–850, 1971. ISSN 0162-1459.

8. R. Satija, J. A. Farrell, D. Gennert, A. F. Schier, and A. Regev. Spatial reconstruction of single-cell gene expression data. *Nature Biotechnology*, 33(5):495–U206, 2015. ISSN 1087-0156.

9. Bosiljka Tasic, Vilas Menon, Thuc Nghi Nguyen, Tae Kyung Kim, Tim Jarsky, Zizhen Yao, Boaz Levi, Lucas T Gray, Staci A Sorensen, Tim Dolbeare, et al. Adult mouse cortical cell taxonomy revealed by single cell transcriptomics. *Nature neuroscience*, 19(2):335–346, 2016.

10. Sandro Vega-Pons and José Ruiz-Shulcloper. A survey of clustering ensemble algorithms. *International Journal of Pattern Recognition and Artificial Intelligence*, 25(03):337–372, 2011.

11. Yue J Wang, Jonathan Schug, Kyoung-Jae Won, Chengyang Liu, Ali Naji, Dana Avrahami, Maria L Golson, and Klaus H Kaestner. Single-cell transcriptomics of the human endocrine pancreas. *Diabetes*, 65(10):3028–3038, 2016.

12. Amit Zeisel, Ana B Muñoz-Manchado, Simone Codeluppi, Peter Lönnerberg, Gioele La Manno, Anna Juréus, Sueli Marques, Hermany Munguba, Liqun He, Christer Betsholtz, et al. Cell types in the mouse cortex and hippocampus revealed by single-cell rna-seq. *Science*, 347(6226): 1138–1142, 2015.

13. X. W. Zhang, C. L. Xu, and N. Yosef. Simulating multiple faceted variability in single cell rna sequencing. *Nature Communications*, 10, 2019. ISSN 2041-1723.

14. L. X. Zhu, J. Lei, B. Devlin, and K. Roeder. A unified statistical framework for single cell and bulk rna sequencing data. *Annals of Applied Statistics*, 12(1):609–632, 2018. ISSN 1932-6157.

15. Lingxue Zhu, Jing Lei, Lambertus Klei, Bernie Devlin, and Kathryn Roeder. Semisoft clustering of single-cell data. *Proceedings of the National Academy of Sciences*, 116(2):466–471, 2019.
